# PlantRegMoD: An integrative and AI-driven multi-omics database for plant regeneration research

**DOI:** 10.64898/2026.09.15.751352

**Authors:** Yue-Min Li, You-Bin Ma, Le-Yue Gui, Yi Tang, Caihong Meng, Yuxin Li, Li Yao, Jixiang Zhang, Siwei Xia, Youhao Peng, Shipeng Song, Zhebin Zeng, Jia-Bao He, Na Zhang, Pei-Xuan Xiao, Yuming Xu, Lei Tan, Akira Iwase, Chunli Chen, Wen-Biao Jiao

**Affiliations:** National Key Laboratory for Germplasm Innovation and Utilization of Horticultural Crops, Huazhong Agricultural University, Wuhan 430070, China; Hubei Hongshan Laboratory, Wuhan 430070, China; Hubei Key Laboratory of Agricultural Bioinformatics, College of Informatics, Huazhong Agricultural University, Wuhan 430070, China; College of Life Science and Technology, Huazhong Agricultural University, Wuhan 430070, China; RIKEN Center for Sustainable Resource Science, Yokohama 230-0045, Japan

**Keywords:** Plant regeneration, Multi-omics, Database, Artificial Intelligence, Retrieval-Augmented Generation

## Abstract

Plant regeneration underpins plant developmental plasticity, tissue culture, genetic transformation and crop improvement. Although numerous related omics datasets have been generated, specialized multi-omics databases for this field are still scarce. Here, we constructed PlantRegMoD, an AI-powered integrated database dedicated to plant regeneration. This platform hosts 20.54 TB standardized multi-omics data from 147 projects across 32 plant species and 2,593 samples. We established a unified hierarchical classification system covering five major categories and nine regeneration models, and curated 236 regeneration genes as well as their 28,190 homologs across 58 representative plant species. It contains extensive transcriptomic resources across all regeneration models, together with 196,423 single cells and over 8.81 million epigenetic peaks to dissect cellular heterogeneity and multi-layered epigenetic regulation. Equipped with nine online omics-related tools and a RAG-based intelligent Q&A system, PlantRegMoD greatly reduces bioinformatic barriers and serves as a robust resource for mechanistic, functional and evolutionary studies of plant regeneration.

## Main Text

Plant regeneration, the remarkable ability of plant cells and tissues to regenerate new organs or entire individuals in response to inductive cues (Duclercq et al., 2011; Ikeuchi et al., 2013), is a fundamental trait underpinning plant survival and adaptation. This process involves extensive cellular reprogramming and the coordinated activation of intricate molecular networks that enable tissue regeneration, *de novo* organogenesis, and stem cell establishment. The efficiency and mechanisms of regeneration are not only crucial for understanding fundamental developmental plasticity (Sugimoto et al., 2011; He et al., 2025) but also directly impact critical applications such as plant tissue culture, genetic transformation, and crop improvement (Zhu et al., 2024). Driven by the rapid advancement of high-throughput technologies, multi-omics studies (genomics, transcriptomics, epigenomics, proteomics, etc.) have begun to unravel the sophisticated molecular choreography governing plant regeneration (Liu et al., 2023; Liao and Wang, 2023; Chen et al., 2024). However, despite the explosion of data generated by these studies, publicly accessible, specialized databases for plant regeneration remain extremely limited. REGENOMICS (Bae et al., 2022), the only currently available regeneration-focused database, merely covers transcriptomic data, lacking epigenetic resources. Furthermore, no existing platform integrates multi-layered omics data with intelligent analytical tools to support user-friendly data mining and hypothesis validation for wet-lab researchers, severely restricting cross-study data integration and mechanistic exploration of plant regeneration.

Here, we constructed PlantRegMoD (Plant Regeneration Multi-omics Database, http://jiao.hzau.edu.cn/prmd/), the first AI-powered integrated multi-omics platform for plant regeneration research. It systematically curates regeneration-related multi-omics data, integrates a standardized regeneration classification framework, multi-dimensional gene features, user-friendly bioinformatic tools, and an intelligent Q&A system for wet-lab researchers **(Figure 1A)**. PlantRegMoD integrates omic data from 147 projects (119 transcriptomic, 6 single-cell omic, and 22 epigenomic) across 32 species, covering 2,593 samples with 20.54 TB of data **(Figure 1B, Table S1-S2)**. All sequencing data were processed with a standardized pipeline to ensure comparability and reliability **(Supplemental Methods)**. This platform comprises seven core modules: *Models, Genes, Transcriptome, Single-cell Transcriptome, Epigenome, Tools, Chatbot* **(Figure S1-S2)**.

**Figure 1.**
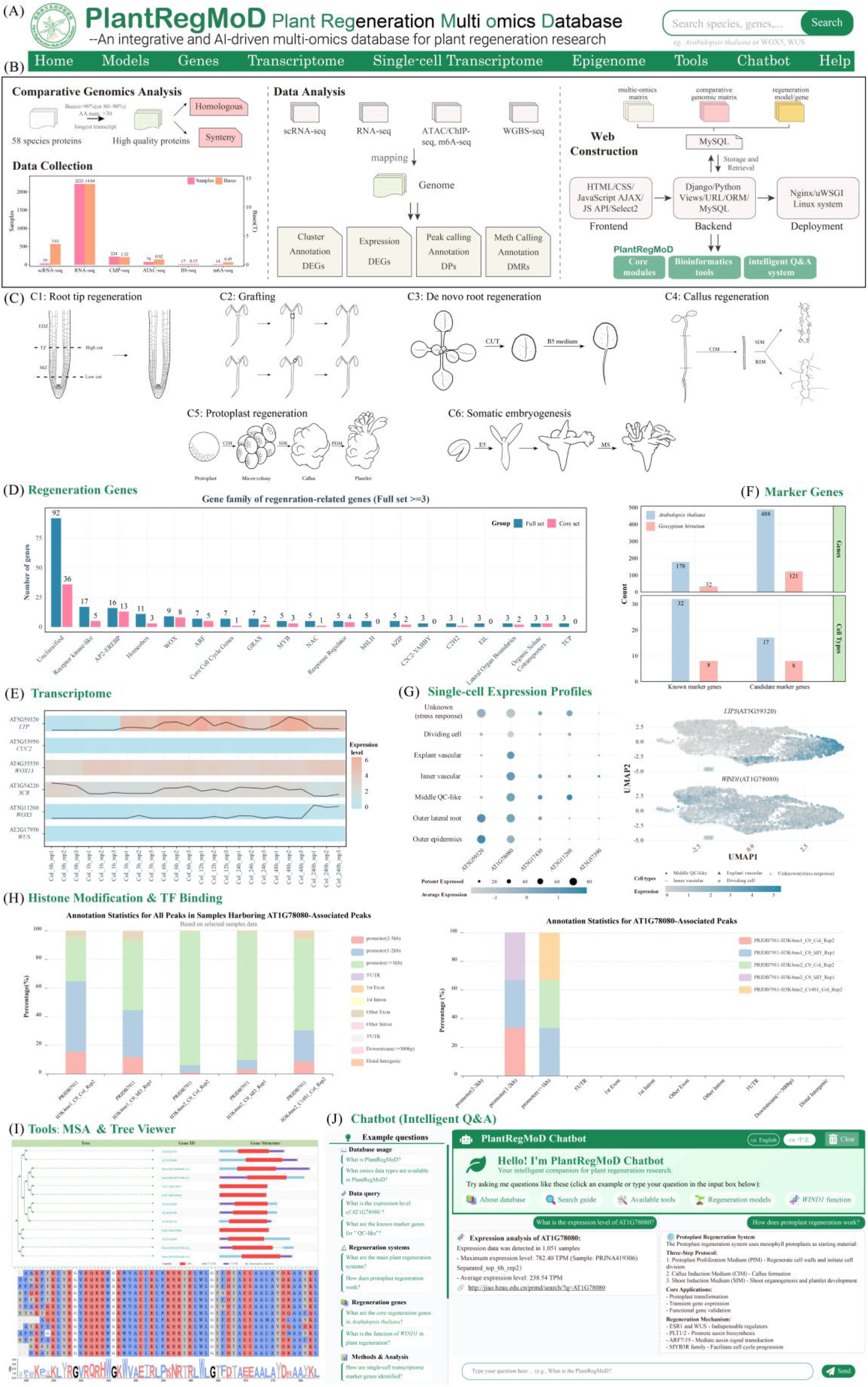
Overview of the Plant Regeneration Multi-omics Database (PlantRegMoD). **(A)** The navigation bar interface of PlantRegMoD. **(B)** The overall workflow for PlantRegMoD construction, including raw data collection, data analysis, and web platform deployment. **(C)** Schematic representation of six categories of plant regeneration systems in the *Models* module. **(D)** Distribution of regeneration-associated gene families (family size > 3; 236 candidate genes (full set) including 102 core genes). **(E)** Visualization of gene expression profiles across samples in the *Transcriptome* module. **(F)** The number of known marker genes and novel candidate marker genes identified in this study for scRNA-seq based cell type annotation. **(G)** Gene expression profiles at single-cell level visualized via dot plot (left) and UMAP (right) in the *Single-cell Transcriptome* module. **(H)** Two types of epigenetic annotation statistics provided by the *Epigenome* module for queried genes or genomic regions: cross-sample global annotation (left) and detailed annotation of corresponding peaks (right). **(I)** Representative usage examples of multiple sequence alignment and phylogenetic tree visualization tools. **(J)** Functional demonstration of the platform’s intelligent Q&A system.

As a highlight of PlantRegMoD, the Models module aims to address the absence of unified classification criteria that has greatly hindered cross-study mechanistic comparison and integration. Based on the previous work (Chen et al., 2024), we further revised and optimized a standardized hierarchical classification system for plant regeneration. This system comprises nine detailed experimental models belonging to six major categories: root tip regeneration, grafting, de novo root regeneration, callus regeneration, protoplast regeneration, and somatic embryogenesis **(Figure 1C, Figure S3)**. For each model, the module delivers information for data summary, pivotal regulatory genes, core molecular regulatory mechanisms, and brief experimental protocols **(Table S3, Figure S3)**. In addition, intuitive schematic diagrams are embedded to visually illustrate the cellular and molecular processes underlying each regeneration model.

To support identifying regeneration genes in non-model plants and characterizing their evolutionary conservation and divergence, the *Genes* module systematically compiles regeneration regulators in *Arabidopsis thaliana*, along with their homologs and genomic synteny across 58 representative plant species covering major plant evolutionary lineages (bryophytes, ferns, gymnosperms, lycophytes, basal angiosperms, monocots and dicots) **(Table S4)**. Taking *A. thaliana* as the well-established reference model, we manually compiled 236 regeneration-associated candidate genes from peer-reviewed studies, among which 102 genes were experimentally validated and defined as core regeneration genes **(Figure 1D, Figure S4**). GO enrichment analysis further verified the high reliability of this gene set **(Figure S4C)**. This module comprises three functional webpages: *Regeneration Genes, Homologs*, and *Micro-synteny*. The *Regeneration Genes* webpage provides comprehensive information for all identified regeneration genes, including gene sequences, functional annotations, gene family classification, literature references, and cross-sample expression and epigenomic profiles **(Figure S5)**. To facilitate cross-species functional extrapolation, the *Homologs* webpage further characterizes homologous genes across the remaining 57 plant species. In total, this module identifies 75 orthologous groups comprising 28,190 regeneration-related genes **(Figure S6)**, and supplies multiple evolutionary analytical results including phylogenetic trees, gene structure diagrams, multiple sequence alignments, and conserved motif maps for cross-species comparative analysis **(Figure S7)**. Furthermore, the *Micro-synteny* webpage enables systematic identification and visualization of pairwise micro-synteny blocks among the 58 plant species via two flexible analytical modes **(Figure S8)**.

Based on the regeneration models and genes curated in this study, the *Transcriptome* module hosts 2,221 transcriptomic datasets tailored for plant regeneration research. It supports gene expression retrieval, differential expression analysis, and multi-dimensional expression visualization for regeneration-associated genes across diverse samples and regeneration models **(Figure S9)**. The module supports three core analytical scenarios: (1) global expression profiling across all or customized sample sets: (2) comparative analysis of expression dynamics across different tissues, treatments, and genetic backgrounds; and (3) temporal expression curve generation for time-course datasets to uncover regulatory cascades **(Figure 1E)**. Additionally, standardized expression matrices (TPM, raw counts, RPKM) are available for download, supporting flexible downstream analysis.

Complementing the bulk transcriptomics, the *Single-cell Transcriptome* module is designed to demonstrate single-cell expression profiles and cellular heterogeneity during generation. This module integrated 39 high-quality scRNA-seq datasets across four regeneration models, covering 196,423 individual cells annotated into 39 cell types (**Table S2**). It consists of three functional pages: *General Statistics, Marker Genes*, and *Single-cell Expression Profiles*. The *General Statistics* page provides standardized quality control and metadata including UMI counts, gene counts, and cell number per cluster. It also presents UMAP and t-SNE dimensionality reduction visualizations, detailed cell annotations, and cluster-specific gene information for each dataset **(Figure S10)**. As the core resource of this module, the *Marker Genes* webpage integrates two complementary sources: 210 experimentally validated marker genes from published studies and 609 newly screened candidate markers identified via rigorous statistical filtering **(Figure 1F, Figure S11, Supplemental Methods)**. Apart from listing these genes, this webpage supports customizable dot plots, violin plots and feature plots for intuitive visualization of cell-type-specific gene expression **(Figure S12)**. The *Single-cell Expression Profiles* page provides both tabular statistics and visualization of the average expression abundance and cell ratio of queried genes across cell types and clusters, to uncover cluster-specific or statistically significant expression signatures during generation **(Figure 1G, Figure S13)**.

Remarkably, PlantRegMod provides the *Epigenome* module, which collates multi-layered transcriptional and epigenomic regulatory profiles derived from all five regeneration models, encompassing a total of 355 samples. This module incorporates 78 ATAC-seq datasets for chromatin accessibility profiling, 224 ChIP-seq datasets for transcription factor binding and histone modification landscapes, 39 BS-seq datasets for DNA methylation mapping, and 14 m^6^A-seq datasets for RNA N6-methyladenosine modification identification **(Table S1)**. It supports flexible data retrieval via gene-based or genomic locus-based queries. For each of 8.81 million annotated peaks **(Table S5)**, this module provides global cross-sample statistics, sample-specific details, and comprehensive peak characterization **(Figure 1H, Figure S14)**. Additionally, for DNA methylation analysis, the module quantifies methylation levels across three canonical sequence contexts (CpG, CHG, and CHH) within gene bodies and promoter regions and characterizes differentially methylated regions to support epigenetic regulatory dissection of regeneration **(Figure S15)**. All epigenomic profiles are visualized through a built-in genome browser, enabling intuitive exploration of regulatory landscapes under diverse plant regeneration contexts **(Figure S14-S15)**.

To enable efficient mining and experimental utilization of the above multi-layered omics resources for wet-lab researchers, the *Tools* module integrates nine user-friendly bioinformatics utilities for sequence analysis, functional enrichment, omics visualization, and experimental design. The sequence analysis toolkits BLAST homology searching, multiple sequence alignment, and phylogenetic visualization **(Figure 1I)**. In addition, this module provides GO and KEGG functional enrichment pipelines, as well as a tool, PlantTF-PK, for transcription factors and protein kinases prediction. The embedded JBrowse genome browser enables flexible interactive visualization and interrogation of multi-omics datasets across different regeneration contexts. Beyond data analysis and visualization, integrated PCR primer and sgRNA design tools further facilitate molecular experimental validation of regeneration-related genes. Meanwhile, a one-stop retrieval function allows comprehensive acquisition of gene sequences, homologous genes, expression profiles, and diverse epigenetic modification profiles **(Figure S16)**.

Notably, to further lower the bioinformatics and interpretative threshold for experimental researchers, we embedded a RAG-based intelligent Q&A system powered by the GLM-4-Flash large language models (Lewis et al., 2021; Lai et al., 2024). Leveraging the curated knowledge base of omics resources and annotated research literature, this system supports natural language interaction for database usage guidance, omics results interpretation, and mechanistic exploration of gene function and plant regeneration regulation **(Figure 1J)**.

In summary, PlantRegMoD integrates standardized regeneration classification systems and high-quality multi-layered omics data, facilitating systematic dissection of the regulatory landscape underlying plant regeneration. More importantly, PlantRegMoD implements versatile bioinformatics tools and an RAG-based intelligent Q&A system, which greatly lowers bioinformatic barriers for wet-lab researchers. PlantRegMoD will receive regular updates of new datasets and functions to provide a valuable community resource for functional, regulatory, and evolutionary studies of plant regeneration.

## Author contributions

Y.-M.L. performed data collection, curation, validation, and analyses, database design and architecture, and wrote the original draft. Y.-B.M. conducted data collection, server deployment, front-end development, data storage, website implementation, and integration of analytical tools and visualizations. L.-Y.G., Y.L., and L.Y. completed the regeneration system classification, schematic illustration, and gene collection and curation. Y.T., C.M., J.Z., S.X., Y.P., S.S., Z.Z., J.-B.H., N.Z., P.-X.X, Y.X., and L.T. contributed to literature curation and database function testing. W.-B.J. conceived and designed the project. W.-B.J. and C.C. supervised the project and revised the manuscript with contributions from A.I.. All authors read and approved the final manuscript.

## Acknowledgments

The authors would like to thank all the research groups who have contributed their valuable multi-omics data (including but not limited to transcriptomic, genomic, and epigenomic data) to the public databases, Masaki Ishikawa (National Institute for Basic Biology), Ling Cui, Kang Xiao, and Jin Lu (Huazhong Agricultural University) for their helpful comments on the database.

## References

Bae, S. H., Noh, Y. S., and Seo, P. J. (2022). REGENOMICS: A web-based application for plant REGENeration-associated transcriptOMICS analyses. Comput. Struct. Biotechnol. J. 20:3234–3247.

Chen, C., Hu, Y., Ikeuchi, M., Jiao, Y., Prasad, K., Su, Y. H., Xiao, J., Xu, L., Yang, W., Zhao, Z., et al. (2024). Plant regeneration in the new era: from molecular mechanisms to biotechnology applications. Sci. China Life Sci. 67:1338–1367.

Duclercq, J., Sangwan-Norreel, B., Catterou, M., and Sangwan, R. S. (2011). De novo shoot organogenesis: from art to science. Trends Plant Sci. 16:597–606.

He, Y., Xu, L., and Liu, Q. (2025). The cellular epigenetic blueprint of plant regeneration. Curr. Opin. Plant Biol. 88:102784.

Ikeuchi, M., Sugimoto, K., and Iwase, A. (2013). Plant Callus: Mechanisms of Induction and Repression. Plant Cell 25:3159–3173.

Lai, H., Wang, B., Zhang, C., Yin, D., Zhang, D., Rojas, D., Feng, G., Zhao, H., Lai, H., Yu, H., et al. (2024). ChatGLM: A Family of Large Language Models from GLM-130B to GLM-4 All Tools Advance Access published 2024, doi:10.48550/ARXIV.2406.12793.

Lewis, P., Perez, E., Piktus, A., Petroni, F., Karpukhin, V., Goyal, N., Küttler, H., Lewis, M., Yih, W., Rocktäschel, T., et al. (2021). Retrieval-Augmented Generation for Knowledge-Intensive NLP Tasks Advance Access published April 12, 2021, doi:10.48550/arXiv.2005.11401.

Liao, R. Y., and Wang, J. W. (2023). Analysis of meristems and plant regeneration at single-cell resolution. Curr. Opin. Plant Biol. 74:102378.

Liu, X., Zhu, K., and Xiao, J. (2023). Recent advances in understanding of the epigenetic regulation of plant regeneration. aBIOTECH 4:31–46.

Sugimoto, K., Gordon, S. P., and Meyerowitz, E. M. (2011). Regeneration in plants and animals: dedifferentiation, transdifferentiation, or just differentiation? Trends Cell Biol. 21:212–218.

Zhu, L., Zhou, L., Li, J., Chen, Z., Wang, M., Li, B., Xu, S., Luo, J., Zeng, T., and Wang, C. (2024). Regeneration of ornamental plants: current status and prospects. Ornam. Plant Res. 4:e022.

