## Supplemental for "PlantRegMoD: An integrative and AI-driven multi-omics database for plant regeneration research"

### **Supplementary Material**

#### **Supplemental Methods**

1. **Data collection**

*Collection of Omic data related to plant regeneration.* A total of 2,452 multi-omics datasets released by previously plant regeneration studies were downloaded from public databases including NCBI, DDBJ, EMBL-EBI, CNGBdb, and NGDC (**Table S1**). Among these resources, unpublished datasets labeled Chen01Gui, Chen02Zhang and Chen03Tang contained *Arabidopsis thaliana* hypocotyl callus RNA-seq data independently generated by us.

*Genome assembly and annotation data*. For 32 species included in this multi-omics dataset resource, their reference genome assembly sequences and gene annotation files were also obtained from NCBI, Ensembl Plants, CNGB (*Areca catechu*), figshare (*Rorippa aquatica*, *Neolamarckia cadamba*), WP-MOD (*Picea abies, Ginkgo biloba*), GCGI (*Gossypium hirsutum*), Phytozome (*Ceratopteris richardii*), BambooBase (*Dendrocalamus latiflorus*), NGDC (*Larix kaempferi*), and TCOD (*Dimocarpus longan*).

*Regulatory genes for plant regeneration*. As *Arabidopsis thaliana* serves as a well-established model for plant regeneration research, many regeneration regulators have been functionally validated in this species. We first integrated stem cell‑related genes from the Plant Stem Cell Database (PSCdb) and performed rigorous literature curation (Wu et al., 2024). After manual verification, 102 genes with close functional relevance to regeneration were retained from 236 initial candidates. All these 102 genes, which have been validated by experiments, are defined as the core regeneration gene set.

2. **Identification of ortholog groups and synteny blocks**

To provide orthologous genes and gene-level synteny information of plant regeneration‑related genes, this database integrated genome assemblies and gene annotations from 58 representative species covering basal angiosperms, core angiosperms, gymnosperms, ferns, and bryophytes (**Table S4**).

All protein sequences were filtered and deduplicated to improve the accuracy of subsequent homology inference and synteny analysis. Briefly, sequences shorter than 30 amino acids were removed, and only the protein sequence translated from the longest transcript of each gene was retained. After data preprocessing, orthologous gene groups were identified using OrthoFinder (v2.5.5) with default parameters (Emms and Kelly, 2019). By aligning these orthologous groups against experimentally verified regeneration regulators in *A. thaliana*, a homologous gene set associated with plant regeneration was constructed for downstream evolutionary analysis. In addition, genome-wide collinearity analysis was conducted using the Python implementation of MCscan under default parameters to identify syntenic blocks harboring regeneration-related genes (Tang et al., 2024).

The online platform’s gene-level micro-synteny visualization module supports two functional modes: *Reference* and *Custom*. In *Reference* mode, micro-synteny blocks of *A. thaliana* regeneration-related genes are visualized across six representative species spanning distinct phylogenetic lineages and regeneration research systems: *Populus trichocarpa* (woody plant regeneration model), *Daucus carota* (somatic embryogenesis model), *Nicotiana tabacum* (classic *in vitro* organogenesis system), *Oryza sativa* (monocot regeneration model), *Phalaenopsis aphrodite* (ornamental plant tissue culture system), and *Amborella trichopoda* (basal angiosperm). Users can also submit target genes from these six species to evaluate their syntenic conservation across lineages. In *Custom* mode, users can select up to six species to compare against a query species for customized micro-synteny analysis, with the platform automatically generating corresponding visualization outputs.

3. **Bulk RNA-seq data analysis**

For bulk RNA-seq data, raw reads were trimmed to remove adapter sequences and low-quality reads using fastp (v0.23.2) (Chen et al., 2018). The resulting cleaned reads were aligned to corresponding reference genomes using Hisat2 (v2.2.1) with default parameters (Kim et al., 2019), and raw gene expression counts were quantified using featureCounts (v2.0.6) (Liao et al., 2014). All downstream analyses were performed in the R environment (v4.3.1). Low-expression genes were filtered using *filterByExpr*, and the resulting expression matrix was subjected to differential expression analysis using both DESeq2 (v1.42.1) and edgeR (v4.0.16) (Robinson et al., 2010; Love et al., 2014). Differentially expressed genes were defined as those with an absolute log_2_-transformed fold change (|log_2_FC|) ≥ 1.5 and a false discovery rate (FDR) ≤ 0.05. For samples with biological replicates, the intersection of results from both methods was adopted to enhance robustness. For samples without replicates, only edgeR results were used.

4. **Single-cell RNA-seq data analysis**

For single-cell RNA-seq (scRNA-seq) data analysis, raw reads were preprocessed and quantified with Cell Ranger (v7.1.0), UMI-tools (v1.1.5), and dnctools (v2.1.2) following the specifications of the respective sequencing platforms (Zheng et al., 2017; Smith et al., 2017). Downstream analyses, including dimensionality reduction, clustering, and cell annotation, were conducted using Seurat (v5.3.0) (Hao et al., 2024). During data initialization, stringent quality filtering was implemented to construct a high-quality primary expression matrix, where genes detected in fewer than five cells (min.cells=5) and cells with fewer than 200 detected genes (min.features=200) were excluded to eliminate sequencing noise, rare sporadic transcripts, empty droplets, and low-quality or dead cells. Cells were further filtered based on an upper limit on the number of genes detected per cell (nfeature.max=5,000) and a mitochondrial gene proportion threshold (mt.percent.threshold=10%). Potential doublets were removed using DoubletFinder (v2.0.4) to obtain a high-quality scRNA-seq expression matrix (McGinnis et al., 2019). For each sample, data normalization and highly variable gene selection were performed using SCTransform. For datasets comprising multiple biological replicates or samples, integration features were first selected across the replicates within the same dataset (SelectIntegrationFeatures, 3,000 features), integration anchors were then identified based on canonical correlation analysis (FindIntegrationAnchors, dims = 1:30), and all samples were finally integrated using IntegrateData; for datasets containing only a single sample, the SCTransform-normalized expression matrix of that sample was used directly for downstream analysis. Based on the resulting expression matrix, principal component analysis and non-linear dimensionality reduction were subsequently performed, followed by cell clustering using the shared nearest neighbor algorithm.

Cluster-specific marker genes were identified via two complementary criteria: (1) genes with adjusted p-value (p_val_adj) < 0.05, average log_2_ fold change (avg_log_2_FC) > 0.25, and expression proportion in the target cluster (pct.1) > 0.25; and (2) genes uniquely enriched in a single cluster, with a detection proportion > 0.1 in the target cluster and < 0.1 in all other clusters. The second criterion enabled the capture of lowly expressed but highly cluster-specific genes that are often missed by conventional screening strategies. Genes satisfying either criterion were integrated to define cluster-specific gene sets, which balanced the robustness of highly expressed markers and the sensitivity of low-expression specific genes, thereby enhancing the accuracy and biological interpretability of cell type annotation. Final cell type identities were determined by matching cluster-specific gene sets against a curated collection of 210 known cell marker genes, which were compiled from six published studies and the PsctH single-cell marker database (Zhang et al., 2021; Zhai and Xu, 2021; Xu et al., 2022; Liu et al., 2022a; Liu et al., 2022b; Zhu et al., 2023; Yin et al., 2024). The full list of these marker genes is available on the PlantRegMoD platform.

In addition, a rigorous multi-step screening pipeline was established to identify novel candidate marker genes based on the scRNA-seq dataset. First, high-specificity preliminary screening was performed to retain genes with a cellular expression rate > 25% in the target cluster and an average expression fraction < 10% across all other clusters to ensure cluster-specific expression and high population coverage. Second, statistical significance validation was conducted by intersecting the pre-screened high-specificity genes with significantly differentially expressed genes. Finally, known marker genes from the curated 210-gene library were excluded from the validated gene set, yielding the final set of novel candidate cell-type marker genes.

5. **Epigenomic sequencing data analysis**

All sequencing raw data were initially processed using fastp (v0.23.2) to remove adapter and low-quality reads. For ATAC-seq and ChIP-seq data, the resulting clean reads were mapped to the corresponding reference genome using Bowtie2 (v2.5.2) (Langmead and Salzberg, 2012), followed by PCR duplicate removal using sambamba (v1.0.1) (Tarasov et al., 2015), and filtering low-quality alignments and reads from organellar genomes using SAMTools (v1.18) (Li et al., 2009). For m^6^A-seq data, clean reads were aligned using Hisat2 (v2.2.1), followed by identical samtools filtering procedures. All three types of epigenome data were subjected to uniform downstream analyses: peak calling with MACS2 (v2.2.9.1) (Zhang et al., 2008), peak annotation with ChIPseeker (v1.38.0) (Yu et al., 2015), and differential peak analysis with DiffBind (v3.12.0).

For whole-genome DNA methylation sequencing data, raw reads were quality-trimmed with fastp prior to alignment. Clean reads were mapped using Bismark (v0.24.2), and PCR duplicates were removed via the built-in *deduplicate_bismark* module, with organelle-derived sequences (mitochondria and chloroplast) discarded simultaneously (Krueger and Andrews, 2011). The BatMeth2 toolkit was then employed for methylation level quantification and annotation (Zhou et al., 2019). Specifically, the *calmeth* module was used to calculate methylation levels, and the *methyGff* module was used to annotate and quantify methylation levels in gene bodies and promoter regions. The *batmeth2_to_bigwig.py* script was executed to generate bigWig files for the visualization of genome-wide methylation landscape. For CpG contexts, *batDMR* was utilized to identify differentially methylated cytosines (DMCs) and differentially methylated regions (DMRs), and the resulting DMRs were finally annotated using ChIPseeker (v1.38.0).

6. **Web-online tools implemented in the platform**

The web platform integrates nine bioinformatics tools in the *Tools* module to support comprehensive bioinformatic analyses for plant regeneration. The embedded JBrowse serves as an interactive genome browser for visualizing genomic features and multi-omics data tracks (Diesh et al., 2023), enabling intuitive browsing of gene structures, peak distributions, methylation patterns, and other genomic annotations. The BLAST tool supports sequence similarity alignment and homology search (Altschul et al., 1990), allowing users to query sequences against the built-in sequence databases of 58 plant species or upload custom sequence libraries. The *MSA Viewer* integrates three mainstream alignment algorithms, including Clustal Omega (default), MAFFT, and MUSCLE (Edgar, 2004; Sievers et al., 2011; Katoh and Standley, 2013), and provides four optional color schemes: Clustal, Hydrophobicity, Charge, and Grayscale. The phylogenetic tree viewer enables tree construction via five algorithms, namely Neighbor-Joining (NJ, default), UPGMA, FastME, Maximum Likelihood (ML), and Maximum Parsimony (MP) (Sokal and Michener, 1958; Fitch, 1971; Felsenstein, 1981; Saitou and Nei, 1987; Lefort et al., 2015).

Functional enrichment analyses of Gene Ontology (GO) and Kyoto Encyclopedia of Genes and Genomes (KEGG) pathways were performed using the enrichGO, enrichKEGG, and universal enricher functions in the clusterProfiler package (Wu et al., 2021). This tool currently supports enrichment analysis for seven plant species, including *Arabidopsis thaliana*, rice, cotton, wheat, tomato, sorghum, and rapeseed, with analysis results visualized as bubble plots or bar charts.

The *PCR Primer* design tool is developed based on the *primer3_core* module of Primer3 (Untergasser et al., 2012). It accepts target gene sequences from the built-in database or user-customized input sequences (uploaded or manually pasted) and generates optimal primer pairs with key quantitative parameters, including primer sequences, melting temperature (Tm), GC content, and amplicon length. The *CRISPR sgRNA Design* tool adopts the CRISPR-Local algorithm to evaluate genome-wide off-target effects and screen high-specificity single-guide RNA (sgRNA) sequences based on local genomic context (Sun et al., 2019). Currently, this tool supports sgRNA design for Cas9 and Cpf1 nuclease systems in *Arabidopsis thaliana* and *Brassica napus*.

7. **Design and implementation of the intelligent Q&A system**

Based on the Retrieval-Augmented Generation (RAG) architecture (Lewis et al., 2021), PlantRegMoD has developed an intelligent question-answering *Chatbot* module tailored to the plant regeneration domain, enabling users to query database contents and regeneration-related scientific questions using natural language. The *Chatbot* employs the Zhipu AI GLM-4-Flash large language model as its natural language understanding and generation engine (Lai et al., 2024), supporting continuous context retention across multi-turn dialogues. The knowledge base is constructed from database help documentation and regeneration system manuals, which are parsed into 208 document chunks. A lightweight keyword-matching-based retrieval strategy is adopted to enable knowledge-enhanced question answering in GPU-free server environments. Additionally, the system incorporates an intent recognition module that parses user queries to identify query intentions and extract key parameters. By invoking existing database API interfaces, it can directly return query results for data types such as gene expression levels, histone modifications, transcription factor binding sites, chromatin accessibility, DNA methylation, m^6^A modifications, and homologous genes, along with page jump links. This enables seamless integration between natural language question answering and database functionalities.

8. **Database implementation**

The database website was run on a Linux system and developed using Nginx (v1.25.3) and uWSGI (v2.0.30). The backend framework was developed using Django (v5.2.3), with MySQL (v8.0.21) employed as the core data storage engine. The frontend interface was implemented with HTML5, CSS3, and JavaScript, and integrated with the Select2 (v4.1.0) library to optimize dropdown menu searching and multi-selection functions. Efficient bidirectional data communication between the web interface and analytical backend was achieved via AJAX and customized RESTful APIs. The embedded R environment (v4.5.1) undertakes all specialized statistical computations and graphical rendering tasks to support diverse downstream analyses.

**References**

**Altschul, S. F., Gish, W., Miller, W., Myers, E. W., and Lipman, D. J.** (1990). Basic local alignment search tool. *Journal of Molecular Biology* **215**:403–410.

**Chen, S., Zhou, Y., Chen, Y., and Gu, J.** (2018). fastp: an ultra-fast all-in-one FASTQ preprocessor. *Bioinformatics* **34**:i884–i890.

**Diesh, C., Stevens, G. J., Xie, P., De Jesus Martinez, T., Hershberg, E. A., Leung, A., Guo, E., Dider, S., Zhang, J., Bridge, C., et al.** (2023). JBrowse 2: a modular genome browser with views of synteny and structural variation. *Genome Biol* **24**:74.

**Edgar, R. C.** (2004). MUSCLE: multiple sequence alignment with high accuracy and high throughput. *Nucleic Acids Res* **32**:1792–1797.

**Emms, D. M., and Kelly, S.** (2019). OrthoFinder: phylogenetic orthology inference for comparative genomics. *Genome Biol* **20**:238.

**Felsenstein, J.** (1981). Evolutionary trees from DNA sequences: A maximum likelihood approach. *J Mol Evol* **17**:368–376.

**Fitch, W. M.** (1971). Toward Defining the Course of Evolution: Minimum Change for a Specific Tree Topology. *Syst Biol* **20**:406–416.

**Hao, Y., Stuart, T., Kowalski, M. H., Choudhary, S., Hoffman, P., Hartman, A., Srivastava, A., Molla, G., Madad, S., Fernandez-Granda, C., et al.** (2024). Dictionary learning for integrative, multimodal and scalable single-cell analysis. *Nat Biotechnol* **42**:293–304.

**Katoh, K., and Standley, D. M.** (2013). MAFFT Multiple Sequence Alignment Software Version 7: Improvements in Performance and Usability. *Mol Biol Evol* **30**:772–780.

**Kim, D., Paggi, J. M., Park, C., Bennett, C., and Salzberg, S. L.** (2019). Graph-based genome alignment and genotyping with HISAT2 and HISAT-genotype. *Nat Biotechnol* **37**:907–915.

**Krueger, F., and Andrews, S. R.** (2011). Bismark: a flexible aligner and methylation caller for Bisulfite-Seq applications. *Bioinformatics* **27**:1571–1572.

**Lai, H., Wang, B., Zhang, C., Yin, D., Zhang, D., Rojas, D., Feng, G., Zhao, H., Lai, H., Yu, H., et al.** (2024). ChatGLM: A Family of Large Language Models from GLM-130B to GLM-4 All Tools Advance Access published July 30, 2024, doi:10.48550/ARXIV.2406.12793.

**Langmead, B., and Salzberg, S. L.** (2012). Fast gapped-read alignment with Bowtie 2. *Nat Methods* **9**:357–359.

**Lefort, V., Desper, R., and Gascuel, O.** (2015). FastME 2.0: A Comprehensive, Accurate, and Fast Distance-Based Phylogeny Inference Program. *Mol Biol Evol* **32**:2798–2800.

**Lewis, P., Perez, E., Piktus, A., Petroni, F., Karpukhin, V., Goyal, N., Küttler, H., Lewis, M., Yih, W., Rocktäschel, T., et al.** (2021). Retrieval-Augmented Generation for Knowledge-Intensive NLP Tasks Advance Access published April 12, 2021, doi:10.48550/arXiv.2005.11401.

**Li, H., Handsaker, B., Wysoker, A., Fennell, T., Ruan, J., Homer, N., Marth, G., Abecasis, G., Durbin, R., and 1000 Genome Project Data Processing Subgroup** (2009). The Sequence Alignment/Map format and SAMtools. *Bioinformatics* **25**:2078–2079.

**Liao, Y., Smyth, G. K., and Shi, W.** (2014). featureCounts: an efficient general purpose program for assigning sequence reads to genomic features. *Bioinformatics* **30**:923–930.

**Liu, G., Li, J., Li, J.-M., Chen, Z., Yuan, P., Chen, R., Yin, R., Liao, Z., Li, X., Gu, Y., et al.** (2022a). Single-cell transcriptome reveals the redifferentiation trajectories of the early stage of de novo shoot regeneration in Arabidopsis thaliana Advance Access published January 2, 2022, doi:10.1101/2022.01.01.474510.

**Liu, W., Zhang, Y., Fang, X., Tran, S., Zhai, N., Yang, Z., Guo, F., Chen, L., Yu, J., Ison, M. S., et al.** (2022b). Transcriptional landscapes of de novo root regeneration from detached Arabidopsis leaves revealed by time-lapse and single-cell RNA sequencing analyses. *Plant Commun* **3**:100306.

**Love, M. I., Huber, W., and Anders, S.** (2014). Moderated estimation of fold change and dispersion for RNA-seq data with DESeq2. *Genome Biol* **15**:550.

**McGinnis, C. S., Murrow, L. M., and Gartner, Z. J.** (2019). DoubletFinder: Doublet Detection in Single-Cell RNA Sequencing Data Using Artificial Nearest Neighbors. *Cell Syst* **8**:329-337.e4.

**Robinson, M. D., McCarthy, D. J., and Smyth, G. K.** (2010). edgeR: a Bioconductor package for differential expression analysis of digital gene expression data. *Bioinformatics* **26**:139–140.

**Saitou, N., and Nei, M.** (1987). The neighbor-joining method: a new method for reconstructing phylogenetic trees. *Mol Biol Evol* **4**:406–425.

**Sievers, F., Wilm, A., Dineen, D., Gibson, T. J., Karplus, K., Li, W., Lopez, R., McWilliam, H., Remmert, M., Söding, J., et al.** (2011). Fast, scalable generation of high‐quality protein multiple sequence alignments using Clustal Omega. *Mol Syst Biol* **7**:MSB201175.

**Smith, T., Heger, A., and Sudbery, I.** (2017). UMI-tools: modeling sequencing errors in Unique Molecular Identifiers to improve quantification accuracy. *Genome Res* **27**:491–499.

**Sokal, R., and Michener, C.** (1958). A statistical method for evaluating systematic relationships. *University of Kansas science bulletin* Advance Access published January 7, 1958.

**Sun, J., Liu, H., Liu, J., Cheng, S., Peng, Y., Zhang, Q., Yan, J., Liu, H.-J., and Chen, L.-L.** (2019). CRISPR-Local: a local single-guide RNA (sgRNA) design tool for non-reference plant genomes. *Bioinformatics* **35**:2501–2503.

**Tang H., Krishnakumar V., Zeng X., Xu Z., Taranto A., Lomas J. S., Zhang Y., Huang Y., Wang Y., Yim W. C., et al.** (2024). JCVI: A versatile toolkit for comparative genomics analysis. *iMeta* **3**:e211.

**Tarasov, A., Vilella, A. J., Cuppen, E., Nijman, I. J., and Prins, P.** (2015). Sambamba: fast processing of NGS alignment formats. *Bioinformatics* **31**:2032–2034.

**Untergasser, A., Cutcutache, I., Koressaar, T., Ye, J., Faircloth, B. C., Remm, M., and Rozen, S. G.** (2012). Primer3—new capabilities and interfaces. *Nucleic Acids Res* **40**:e115.

**Wu, T., Hu, E., Xu, S., Chen, M., Guo, P., Dai, Z., Feng, T., Zhou, L., Tang, W., Zhan, L., et al.** (2021). clusterProfiler 4.0: A universal enrichment tool for interpreting omics data. *Innovation (Camb)* **2**:100141.

**Wu, X., Yuan, Y., Zhou, S., Wang, Z., Li, H., Wu, W., Lei, Z., Liu, S., Qi, K., Yin, H., et al.** (2024). Plant Stem Cell Informatics Database (PSCIdb): A comprehensive computational platform for identifying and analyzing genes related to plant stem cells. *Plant Communications* **5**:100818.

**Xu, Z., Wang, Q., Zhu, X., Wang, G., Qin, Y., Ding, F., Tu, L., Daniell, H., Zhang, X., and Jin, S.** (2022). Plant Single Cell Transcriptome Hub (PsctH): an integrated online tool to explore the plant single-cell transcriptome landscape. *Plant Biotechnology Journal* **20**:10–12.

**Yin, R., Chen, R., Xia, K., and Xu, X.** (2024). A single-cell transcriptome atlas reveals the trajectory of early cell fate transition during callus induction in Arabidopsis. *Plant Commun* **5**:100941.

**Yu, G., Wang, L. G., and He, Q. Y.** (2015). ChIPseeker: an R/Bioconductor package for ChIP peak annotation, comparison and visualization. *Bioinformatics* **31**:2382–2383.

**Zhai, N., and Xu, L.** (2021). Pluripotency acquisition in the middle cell layer of callus is required for organ regeneration. *Nat. Plants* **7**:1453–1460.

**Zhang, Y., Liu, T., Meyer, C. A., Eeckhoute, J., Johnson, D. S., Bernstein, B. E., Nusbaum, C., Myers, R. M., Brown, M., Li, W., et al.** (2008). Model-based analysis of ChIP-Seq (MACS). *Genome Biol* **9**:R137.

**Zhang, T. Q., Chen, Y., and Wang, J. W.** (2021). A single-cell analysis of the Arabidopsis vegetative shoot apex. *Developmental Cell* **56**:1056-1074.e8.

**Zheng, G. X. Y., Terry, J. M., Belgrader, P., Ryvkin, P., Bent, Z. W., Wilson, R., Ziraldo, S. B., Wheeler, T. D., McDermott, G. P., Zhu, J., et al.** (2017). Massively parallel digital transcriptional profiling of single cells. *Nat Commun* **8**:14049.

**Zhou, Q., Lim, J. Q., Sung, W. K., and Li, G.** (2019). An integrated package for bisulfite DNA methylation data analysis with Indel-sensitive mapping. *BMC Bioinformatics* **20**:47.

**Zhu, X., Xu, Z., Wang, G., Cong, Y., Yu, L., Jia, R., Qin, Y., Zhang, G., Li, B., Yuan, D., et al.** (2023). Single-cell resolution analysis reveals the preparation for reprogramming the fate of stem cell niche in cotton lateral meristem. *Genome Biol* **24**:194.

#### **Supplemental Tables**

**Supplemental Tables S1-S5**

**Table S1 The summary of omic data collected in this study.**

| **Omics** | **Projects** | **Datasets/Samples** | **Total Bases(T)** |
| --- | --- | --- | --- |
| scRNA-seq | 6 | 39 | 3.61 |
| RNA-seq | 119 | 2221 | 14.04 |
| ChIP-seq | 13 | 224 | 1.32 |
| ATAC-seq | 4 | 78 | 0.97 |
| BS-seq | 3 | 17 | 0.15 |
| m^6^A-seq | 2 | 14 | 0.45 |
| Total | 147 | 2,593 | 20.54 |

**Table S2 Overview of scRNA-seq datasets used in this study.**

| **Projects** | **Number of cells** | **Detected genes** | **Number of cell types** | **Number of clusters** |
| --- | --- | --- | --- | --- |
| PRJNA895970 | 28,996 | 46,038 | 8 | 10 |
| PRJNA895968 | 30,879 | 45,584 | 7 | 8 |
| CNP0004389 | 30,344 | 24,167 | 17 | 24 |
| PRJNA613684 | 6,327 | 20,778 | 10 | 18 |
| CNP0002343 | 94,593 | 20,784 | 7 | 9 |
| PRJNA659737 | 5,284 | 21,187 | 7 | 11 |

**Table S3 Datasets summary in this study by *Models*.**

| **Model** | **Sub-model** | **Projects** | **Samples** |
| --- | --- | --- | --- |
| Category 1 Root tip regeneration | Model1 | 1 | 12 |
| Category 2 Grafting | Model2 | 3 | 166 |
| Category 3 De novo root regeneration | Model3 | 13 | 309 |
| Category 4 Callus regeneration | Model4.1 | 22 | 283 |
|  | Model4.2 | 18 | 408 |
|  | Model4.3 | 15 | 282 |
| Category 5 Protoplast regeneration | Model5 | 2 | 65 |
| Category 6 Somatic embryogenesis | Model6.1 | 26 | 497 |
|  | Model6.2 | 28 | 312 |
|  | Model6.1/6.2 | 5 | 30 |
| Other | Other | 12 | 221 |
| Uncertain | Uncertain | 2 | 8 |

**Note:** "Other" refers to regeneration systems that do not fall within the six major categories defined above. "Uncertain" refers to datasets that could not be accurately classified due to incomplete descriptions or unconfirmed factors such as explant origin. "Model6.1/Model6.2" indicates datasets that cover both Model6.1 and Model6.2 developmental stages, spanning from embryogenic initiation through subsequent shared developmental stages.

**Table S4 Genomic data used in the *Genes* module.**

| **Clade** | **Species** | **Busco** | **Total proteins** | **Filtered proteins** | **Protein source** | **Version** |
| --- | --- | --- | --- | --- | --- | --- |
| Basal angiosperms | *Amborella trichopoda* | C:98.0%[S:48.0%,D:49.9%],F:0.9%,M:1.1%,n:1614 | 31,494 | 17,105 | NCBI | AMTR1.0 |
| Basal angiosperms | *Nymphaea colorata* | C:97.6%[S:56.6%,D:41.0%],F:0.1%,M:2.4%,n:1614 | 34,619 | 20,487 | NCBI | ASM883128v2 |
| Basal angiosperms | *Victoria cruziana* | C:94.4%[S:47.0%,D:47.3%],F:1.1%,M:4.5%,n:1614 | 48,823 | 26,856 | NCBI | VB03 |
| Bryophytes | *Marchantia polymorpha* | C:87.5%[S:68.8%,D:18.6%],F:1.2%,M:11.3%,n:1614 | 24,674 | 19,284 | EnsemblPlants | Marchanta_polymorpha_v1 |
| Bryophytes | *Physcomitrium patens* | C:88.7%[S:1.0%,D:87.7%],F:1.3%,M:10.0%,n:1614 | 86,669 | 32,009 | EnsemblPlants | Phypa_V3 |
| Dicots | *Actinidia chinensis* | C:94.4%[S:73.6%,D:20.8%],F:2.5%,M:3.0%,n:1614 | 33,115 | 33,042 | EnsemblPlants | Red5_PS1_1.69.0 |
| Dicots | *Arabidopsis thaliana* | C:97.6%[S:6.7%,D:90.9%],F:0.5%,M:1.9%,n:1614 | 48,321 | 27,573 | EnsemblPlants | TAIR10 |
| Dicots | *Brassica napus* | C:97.6%[S:6.7%,D:90.9%],F:0.5%,M:1.9%,n:1614 | 101,040 | 99,741 | EnsemblPlants | AST_PRJEB5043_v1 |
| Dicots | *Brassica rapa* | C:96.7%[S:82.9%,D:13.8%],F:1.6%,M:1.7%,n:1614 | 41,025 | 41,018 | EnsemblPlants | Brapa_1.0 |
| Dicots | *Camellia sinensis* | C:96.5%[S:47.3%,D:49.2%],F:1.7%,M:1.7%,n:1614 | 76,698 | 50,837 | NCBI | AHAU_CSS_1 |
| Dicots | *Capsicum annuum* | C:98.9%[S:56.6%,D:42.3%],F:0.6%,M:0.4%,n:1614 | 51,555 | 32,201 | NCBI | UCD10Xv1 |
| Dicots | *Citrullus lanatus* | C:92.8%[S:91.3%,D:1.5%],F:3.9%,M:3.3%,n:1614 | 22,541 | 22,541 | EnsemblPlants | Cla97_v1 |
| Dicots | *Citrus clementina* | C:93.6%[S:70.1%,D:23.5%],F:4.7%,M:1.7%,n:1614 | 34,557 | 24,950 | EnsemblPlants | Citrus_clementina_v1.0 |
| Dicots | *Citrus hindsii* | C:96.3%[S:52.7%,D:43.6%],F:1.6%,M:2.0%,n:1614 | 52,686 | 32,257 | CPBD | GJ.v1.0 |
| Dicots | *Citrus sinensis* | C:99.7%[S:57.0%,D:42.7%],F:0.1%,M:0.2%,n:1614 | 40,427 | 23,555 | NCBI | DVS_A1.0 |
| Dicots | *Cucumis melo* | C:98.9%[S:56.8%,D:42.2%],F:0.4%,M:0.7%,n:1614 | 35,817 | 20,741 | NCBI | USDA_Cmelo_AY_1.0 |
| Dicots | *Cucumis sativus* | C:91.3%[S:90.6%,D:0.7%],F:5.4%,M:3.3%,n:1614 | 23,780 | 23,778 | EnsemblPlants | ASM407v2 |
| Dicots | *Cucurbita pepo* | C:98.8%[S:54.2%,D:44.5%],F:0.6%,M:0.7%,n:1614 | 43,464 | 29,279 | NCBI | ASM280686v2 |
| Dicots | *Daucus carota* | C:99.4%[S:65.2%,D:34.3%],F:0.1%,M:0.4%,n:1614 | 49,321 | 33,078 | NCBI | DH1v3.0 |
| Dicots | *Dimocarpus longan* | C:97.9%[S:87.4%,D:10.5%],F:1.1%,M:1.1%,n:1614 | 42,155 | 40,353 | TCOD | GWHBDNI00000000 |
| Dicots | *Eucalyptus grandis* | C:92.3%[S:62.9%,D:29.4%],F:4.7%,M:3.0%,n:1614 | 46,920 | 36,621 | EnsemblPlants | Egrandis1_0 |
| Dicots | *Fragaria vesca* | C:96.3%[S:92.6%,D:3.7%],F:0.9%,M:2.8%,n:1614 | 23,319 | 22,379 | NCBI | FraVesHawaii_1.0 |
| Dicots | *Glycine max* | C:99.3%[S:24.7%,D:74.5%],F:0.4%,M:0.4%,n:1614 | 88,412 | 55,890 | EnsemblPlants | Glycine_max_v2.1 |
| Dicots | *Gossypium hirsutum* | C:99.4%[S:10.2%,D:89.2%],F:0.3%,M:0.2%,n:1614 | 115,835 | 70,198 | GCCI | Gh_HAU_v1.1 |
| Dicots | *Helianthus annuus* | C:97.0%[S:86.0%,D:11.0%],F:1.5%,M:1.5%,n:1614 | 70,864 | 70,864 | EnsemblPlants | HanXRQr2.0-SUNRISE |
| Dicots | *Liquidambar formosana* | C:83.1%[S:72.4%,D:10.8%],F:11.4%,M:5.5%,n:1614 | 28,041 | 28,040 | NCBI | ZJU_Lfor_1.0 |
| Dicots | *Malus domestica* | C:93.9%[S:65.2%,D:28.6%],F:3.3%,M:2.9%,n:1614 | 40,624 | 40,624 | EnsemblPlants | ASM211411v1 |
| Dicots | *Manihot esculenta* | C:99.6%[S:52.4%,D:47.3%],F:0.3%,M:0.1%,n:1614 | 59,149 | 32,800 | EnsemblPlants | M.esculenta_v8 |
| Dicots | *Medicago truncatula* | C:97.3%[S:92.6%,D:4.7%],F:2.0%,M:0.7%,n:1614 | 44,450 | 44,450 | EnsemblPlants | MtrunA17r5.0_ANR |
| Dicots | *Nicotiana tabacum* | C:98.1%[S:13.9%,D:84.2%],F:1.7%,M:0.2%,n:1614 | 84,255 | 60,380 | NCBI | ASM71507v2 |
| Dicots | *Populus trichocarpa* | C:99.3%[S:60.5%,D:38.7%],F:0.2%,M:0.6%,n:1614 | 52,400 | 34,681 | EnsemblPlants | Pop_tri_v4 |
| Dicots | *Prunus persica* | C:99.6%[S:69.2%,D:30.4%],F:0.1%,M:0.3%,n:1614 | 32,595 | 23,134 | NCBI | Prunus_persica_NCBIv2 |
| Dicots | *Pyrus communis* | C:98.8%[S:49.9%,D:48.8%],F:0.2%,M:1.1%,n:1614 | 43,421 | 34,369 | NCBI | drPyrComm1.1 |
| Dicots | *Solanum lycopersicum* | C:94.2%[S:75.5%,D:18.7%],F:2.3%,M:3.5%,n:1614 | 43,752 | 36,648 | SGN/TGG | SL5.0 |
| Dicots | *Solanum tuberosum* | C:98.2%[S:95.5%,D:2.7%],F:1.1%,M:0.7%,n:1614 | 40,957 | 40,950 | bioinformaticslab-dm8 | DM8 |
| Dicots | *Theobroma cacao* | C:99.4%[S:55.3%,D:44.1%],F:0.2%,M:0.4%,n:1614 | 44,186 | 29,188 | EnsemblPlants | Theobroma_cacao_20110822 |
| Dicots | *Vitis vinifera* | C:98.9%[S:79.2%,D:19.7%],F:0.7%,M:0.4%,n:1614 | 41,097 | 35,134 | EnsemblPlants | PN40024.v4 |
| Ferns | *Ceratopteris richardii* | C:84.0%[S:33.6%,D:50.4%],F:4.8%,M:11.2%,n:1614 | 75,253 | 36,850 | Phytozome | Crichardii_676_v2.1 |
| Ferns | *Salvinia cucullata* | C:85.9%[S:81.0%,D:5.0%],F:1.6%,M:12.5%,n:1614 | 22,606 | 22,565 | fernbase | v2 |
| Gymnosperms | *Cycas panzhihuaensis* | C:90.1%[S:84.5%,D:5.6%],F:2.7%,M:7.1%,n:1614 | 32,353 | 32,353 | WP-MOD | - |
| Gymnosperms | *Gnetum montanum* | C:83.3%[S:79.2%,D:4.1%],F:4.5%,M:12.3%,n:1614 | 27,354 | 27,354 | CNGB | - |
| Gymnosperms | *Metasequoia glyptostroboides* | C:81.0%[S:76.3%,D:4.6%],F:6.6%,M:12.5%,n:1614 | 32,174 | 32,174 | NGDC | GWHCBJF00000000 |
| Gymnosperms | *Taxus wallichiana* | C:84.1%[S:77.1%,D:6.9%],F:8.1%,M:7.8%,n:1614 | 44,035 | 44,006 | CNGB | - |
| Lycophytes | *Selaginella moellendorffii* | C:87.2%[S:4.8%,D:82.3%],F:1.8%,M:11.0%,n:1614 | 45,247 | 33,936 | NCBI | GCF_000143415.4_v1.0 |
| Monocots | *Ananas comosus* | C:99.1%[S:56.4%,D:42.7%],F:0.6%,M:0.2%,n:1614 | 35,775 | 22,250 | NCBI | ASM154086v1 |
| Monocots | *Apostasia shenzhenica* | C:81.5%[S:80.0%,D:1.4%],F:8.4%,M:10.2%,n:1614 | 21,743 | 21,743 | NCBI | ASM278626v1 |
| Monocots | *Areca catechu* | C:87.0%[S:82.5%,D:4.5%],F:6.2%,M:6.8%,n:1614 | 31,569 | 31,569 | CNGB | v1.0 |
| Monocots | *Dendrobium catenatum* | C:93.2%[S:54.1%,D:39.1%],F:3.6%,M:3.2%,n:1614 | 34,389 | 22,642 | NCBI | ASM160598v2 |
| Monocots | *Dendrocalamus latiflorus* | C:98.1%[S:5.9%,D:92.3%],F:0.9%,M:1.0%,n:1614 | 135,231 | 112,636 | BambooBase | - |
| Monocots | *Elaeis guineensis* | C:97.5%[S:48.9%,D:48.6%],F:0.2%,M:2.3%,n:1614 | 48,771 | 28,076 | NCBI | EG11 |
| Monocots | *Hordeum vulgare* | C:93.9%[S:83.8%,D:10.1%],F:0.8%,M:5.3%,n:1614 | 37,961 | 35,825 | EnsemblPlants | MorexV3 |
| Monocots | *Oryza sativa* | C:99.1%[S:85.3%,D:13.8%],F:0.5%,M:0.4%,n:1614 | 42,853 | 35,717 | RAP | IRGSP-1.0 |
| Monocots | *Panicum miliaceum* | C:94.8%[S:35.1%,D:59.7%],F:2.2%,M:3.0%,n:1614 | 55,964 | 55,964 | milletdb | AN00000390 |
| Monocots | *Phalaenopsis aphrodite* | C:94.1%[S:92.4%,D:1.7%],F:2.4%,M:3.5%,n:1614 | 28,910 | 28,903 | OrchidMD | - |
| Monocots | *Phalaenopsis equestris* | C:94.3%[S:57.2%,D:37.1%],F:2.6%,M:3.1%,n:1614 | 29,894 | 20,153 | NCBI | ASM126359v1 |
| Monocots | *Sorghum bicolor* | C:98.3%[S:70.1%,D:28.2%],F:0.9%,M:0.8%,n:1614 | 47,110 | 34,116 | EnsemblPlants | Sorghum_bicolor_NCBIv3 |
| Monocots | *Triticum aestivum* | C:99.3%[S:0.9%,D:98.3%],F:0.0%,M:0.7%,n:1614 | 133,346 | 107,544 | EnsemblPlants | IWGSC |
| Monocots | *Zea mays* | C:96.3%[S:40.6%,D:55.7%],F:0.9%,M:2.8%,n:1614 | 72,539 | 39,756 | EnsemblPlants | Zm-B73-REFERENCE-NAM-5.0 |

Note: "-" indicates version information is unavailable.

**Table S5 Summary of epigenomic data analysis results in the *Epigenome* module.**

| **Data** | **Peaks** | **Genes** | **Differential peaks or methylation regions** | **Genes related to differential peaks or methylation regions** |
| --- | --- | --- | --- | --- |
| ChIP-seq | 4,626,736 | 140,314 | 3,265,725 | 110,566 |
| ATAC-seq | 4,096,034 | 127,601 | 1,535,779 | 109,585 |
| m^6^A-seq | 83,917 | 24,285 | 7,751 | 3,178 |
| BS-seq | - | - | 42,957 | 16,240 |

#### **Supplemental Figures**

**Supplemental Figures S1-S16**

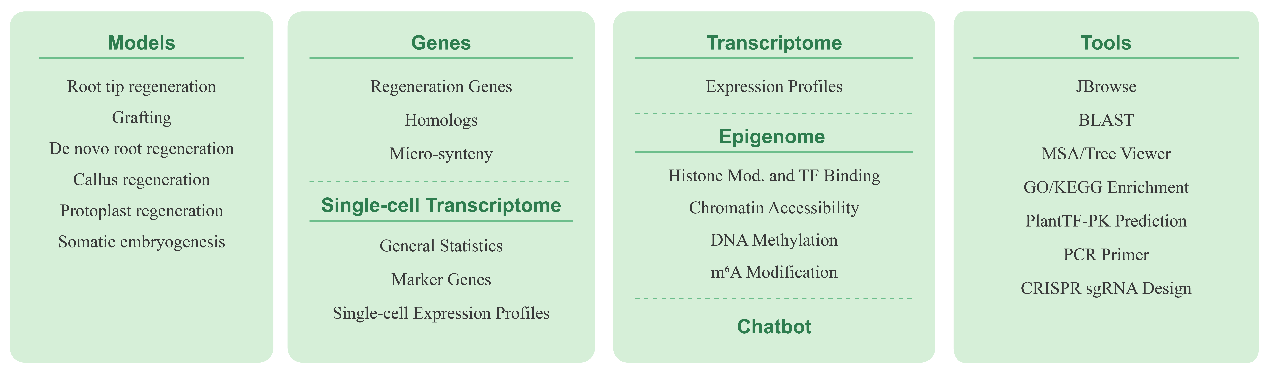

**Figure S1. The core modules in PlantRegMoD.**

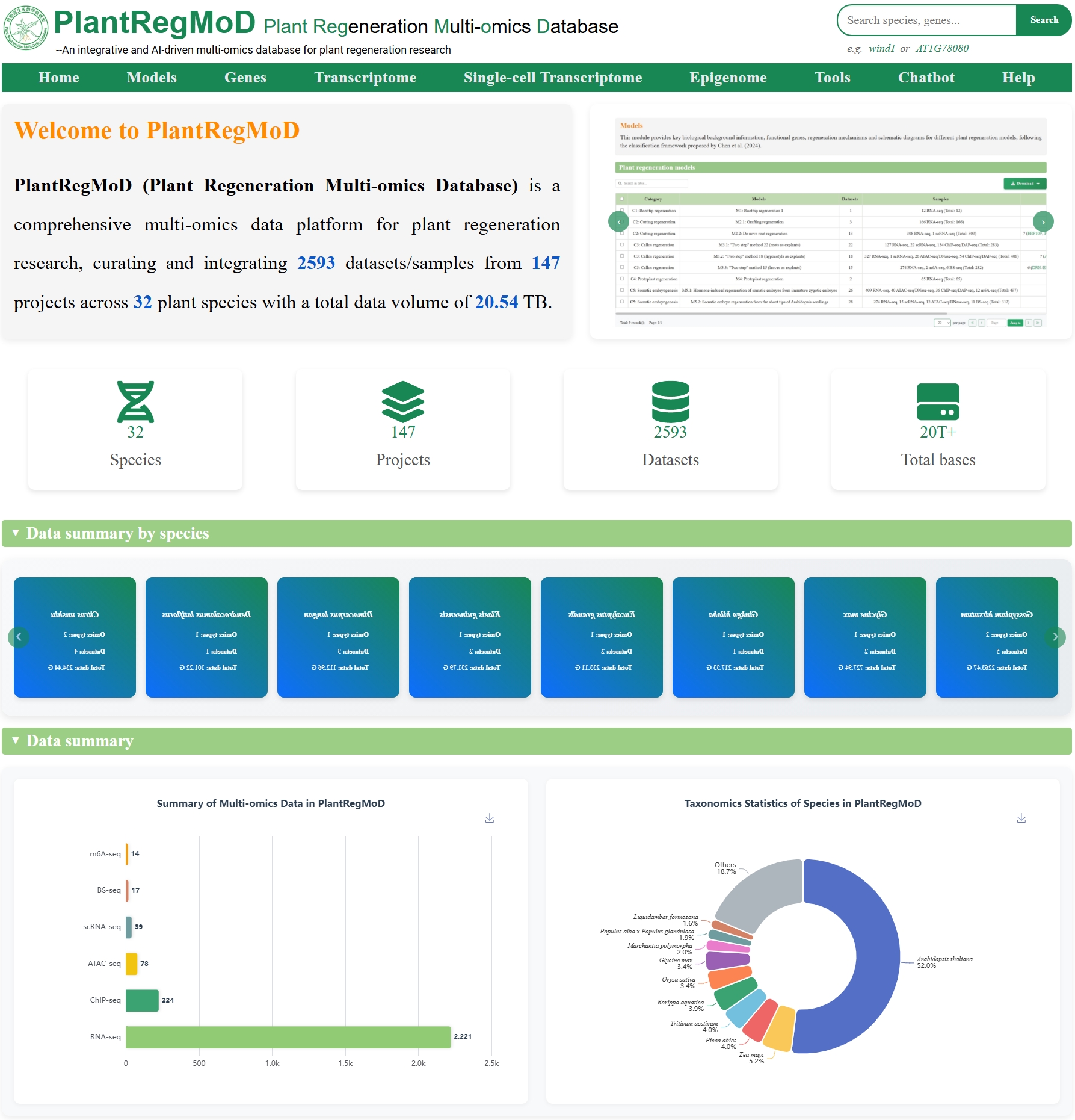

**Figure S2. Screenshots showing the *Home* page in PlantRegMoD.**

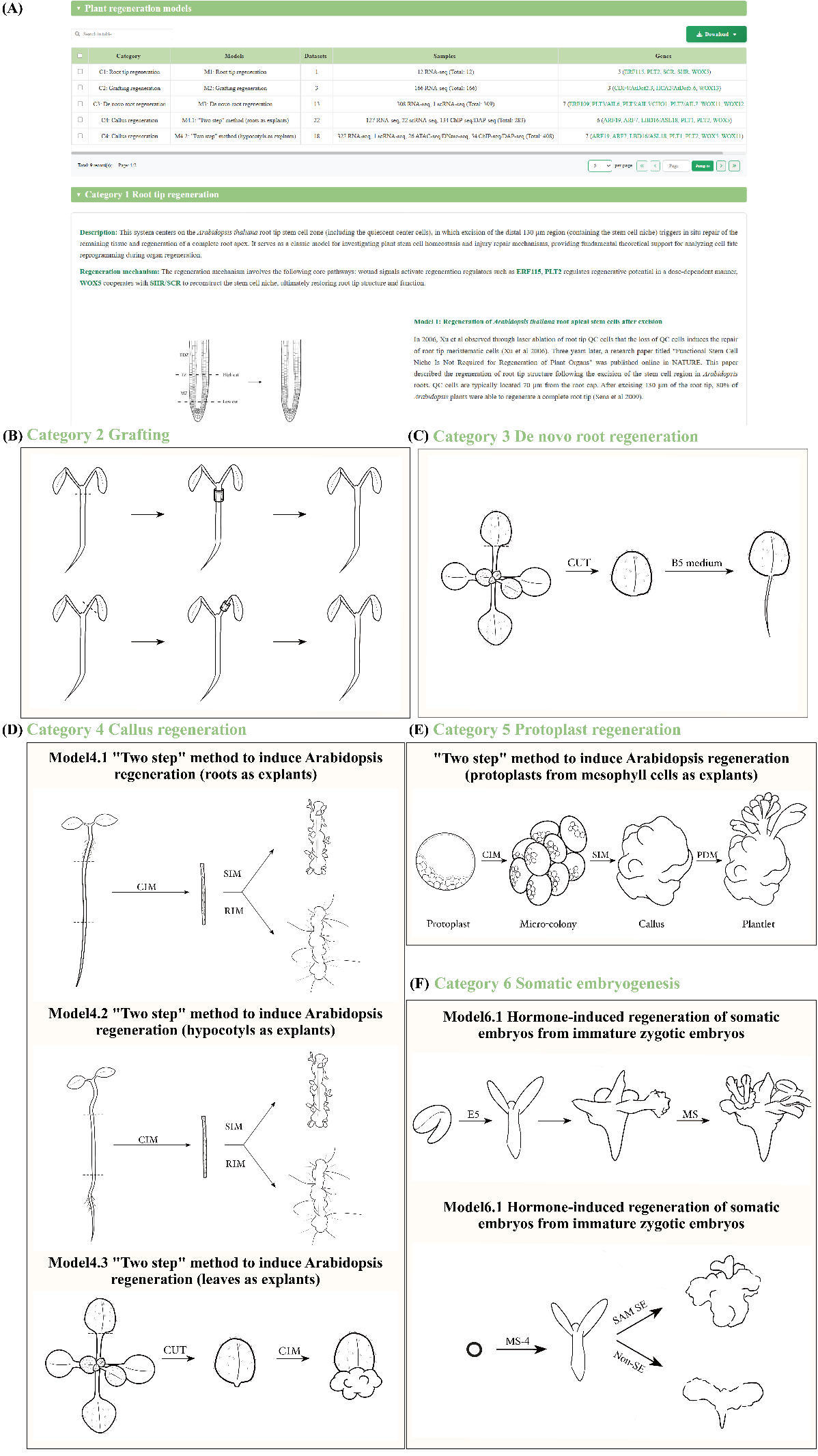

**Figure S3 Systematic classification of other plant regeneration systems in the *Models* module.**

(A) Screenshots showing the plant regeneration models in *Models*. (B)-(F) Schematic diagrams for the different plant regeneration models.

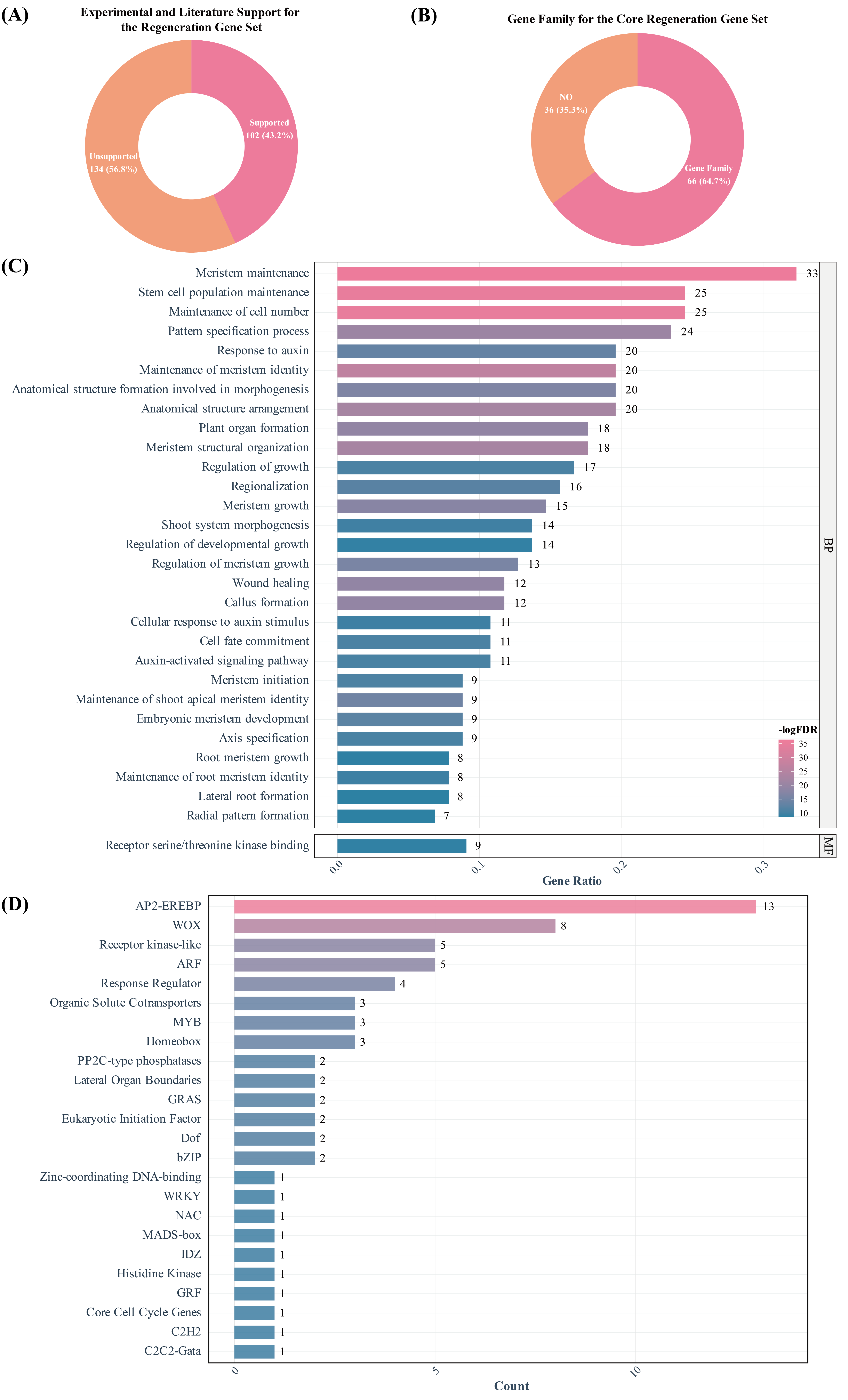

**Figure S4 Overview of the core regeneration genes.**

(A) Proportion of the core regeneration gene with experiment validation or only literature support. (B) Proportional of the core regeneration genes with or without gene family information. (C) GO enrichment analysis of the core regeneration genes.

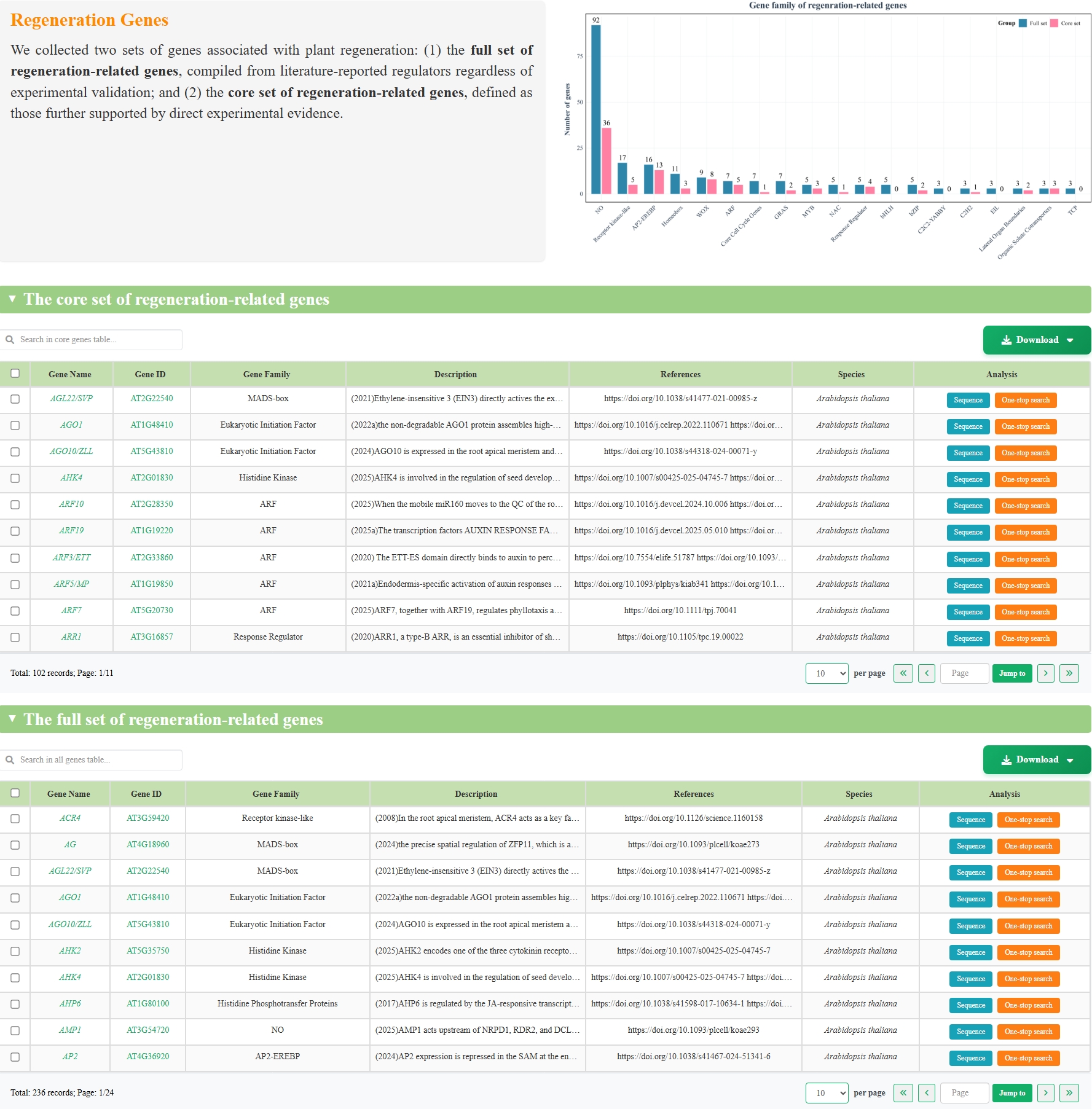

**Figure S5 Screenshots showing the *Regeneration Genes* webpage in *Genes* module.**

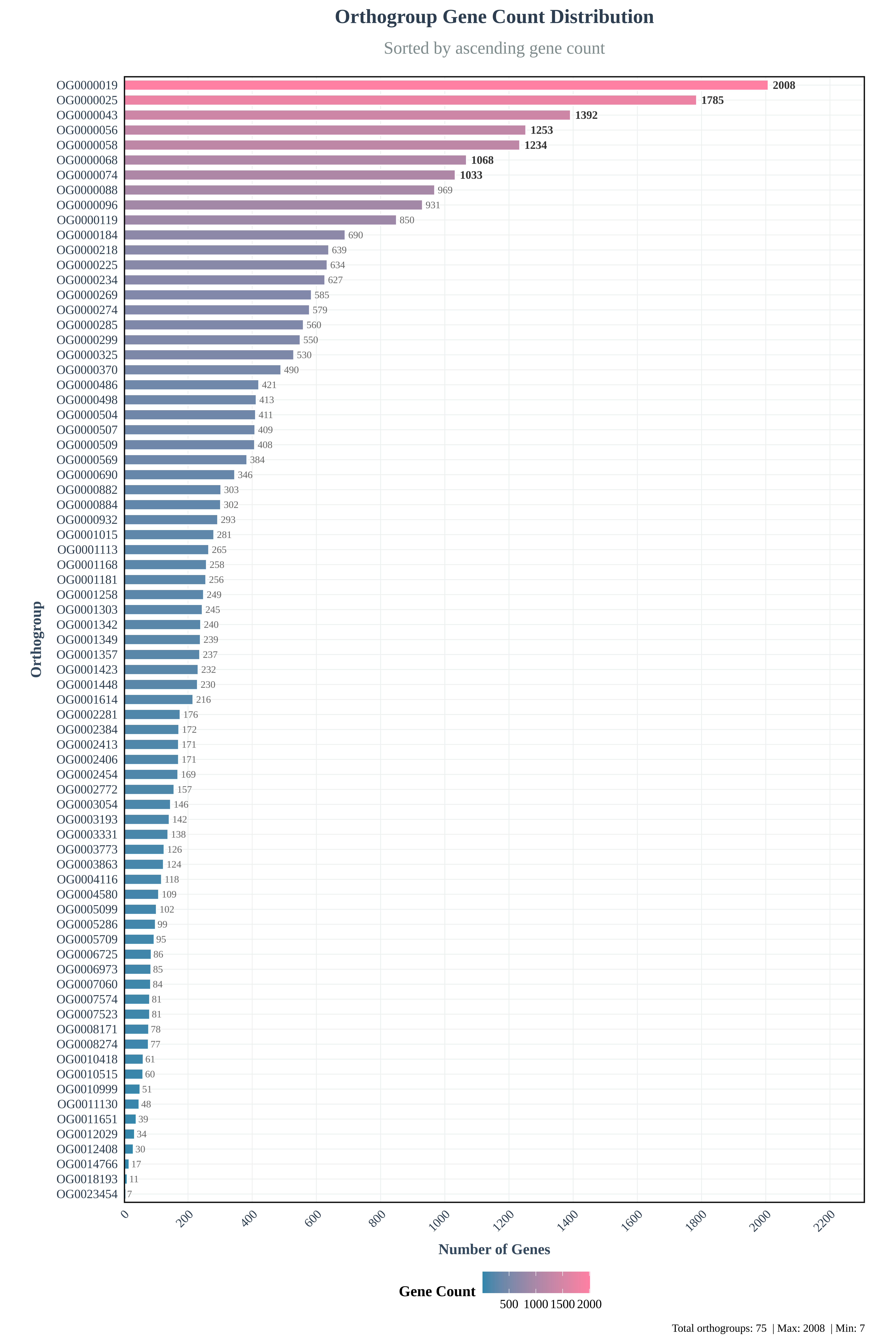

**Figure S6 Gene count of ortholog groups with core regeneration gene.**

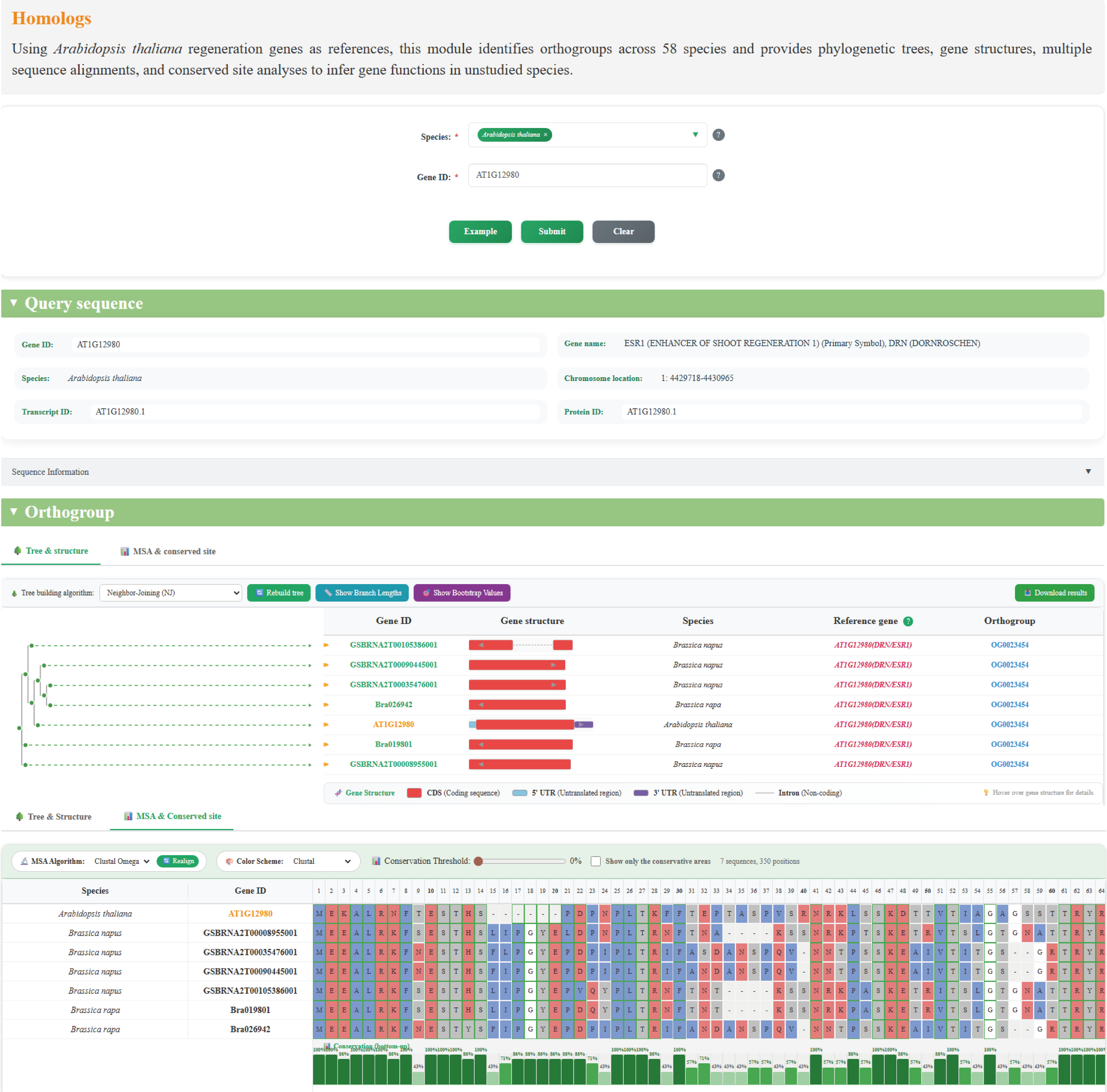

**Figure S7. Screenshot showing the homologous group of the *ESR1* (AT1G12980*)* retrieved in the *Homologs* webpage.**

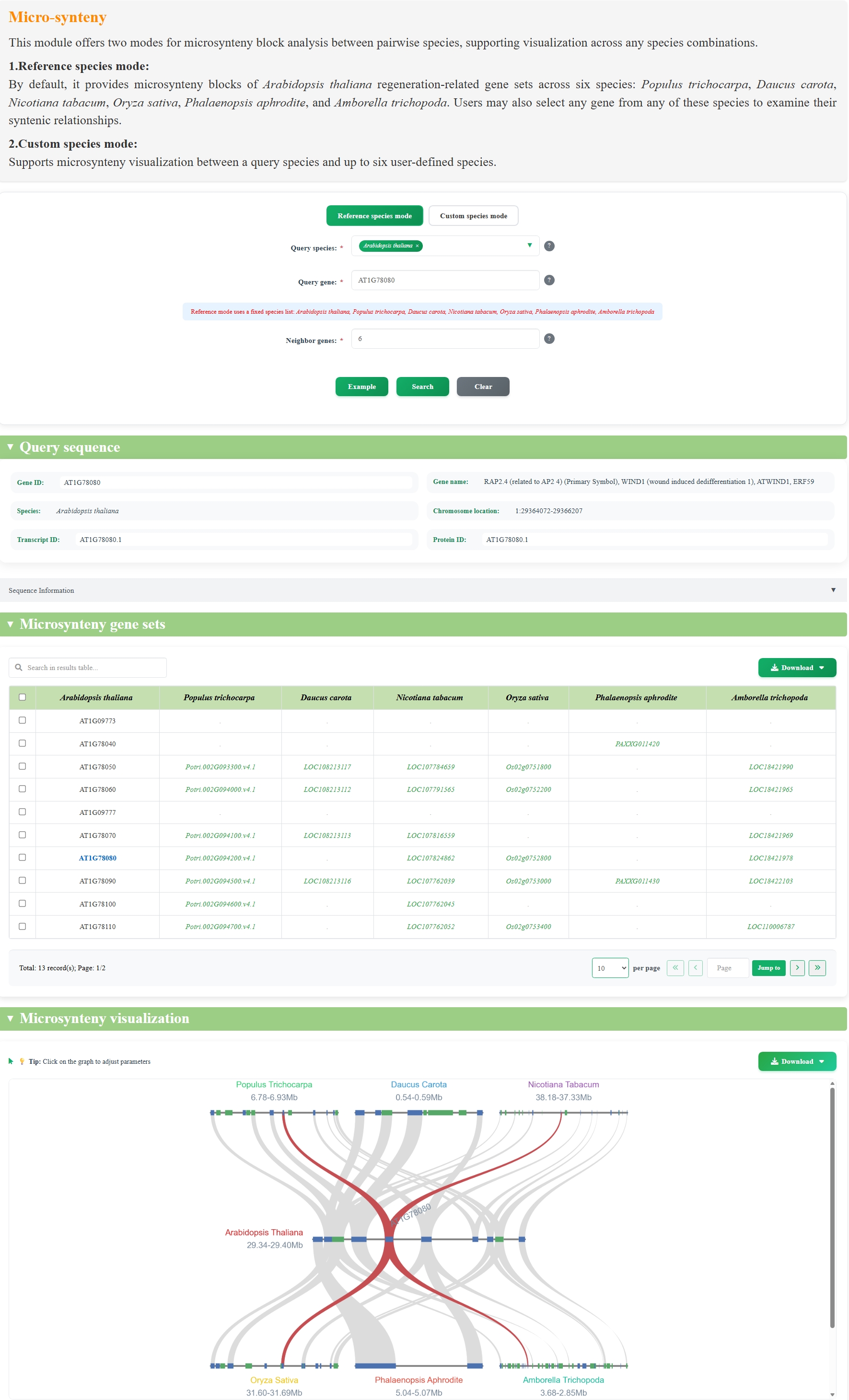

**Figure S8. Screenshots showing the micro-synteny relationship for *WIND1* generated using the *Reference Species* mode in the *Micro-synteny* webpage.**

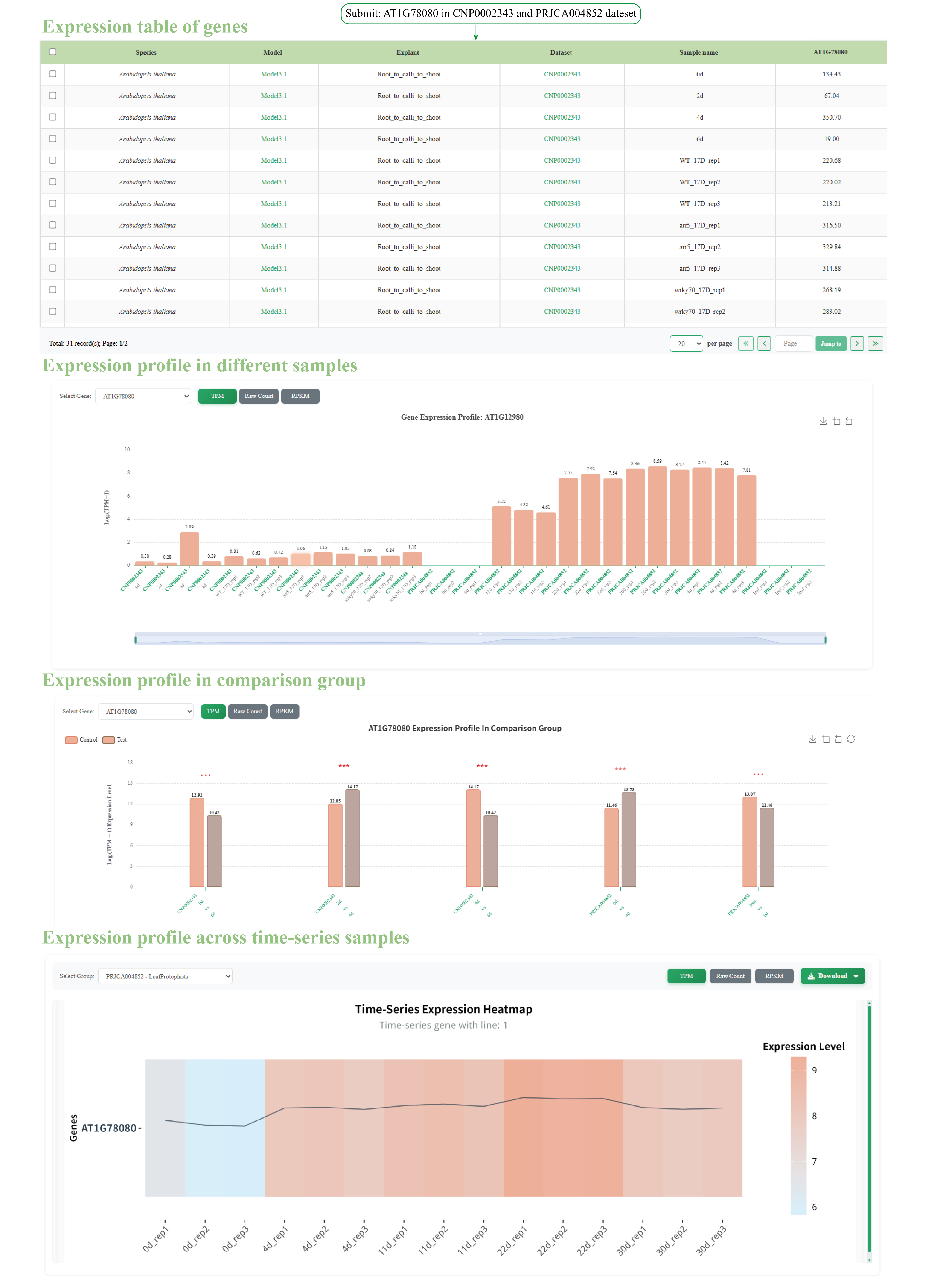

**Figure S9 Screenshots showing the *Transcriptome* module in PlantRegMoD.**

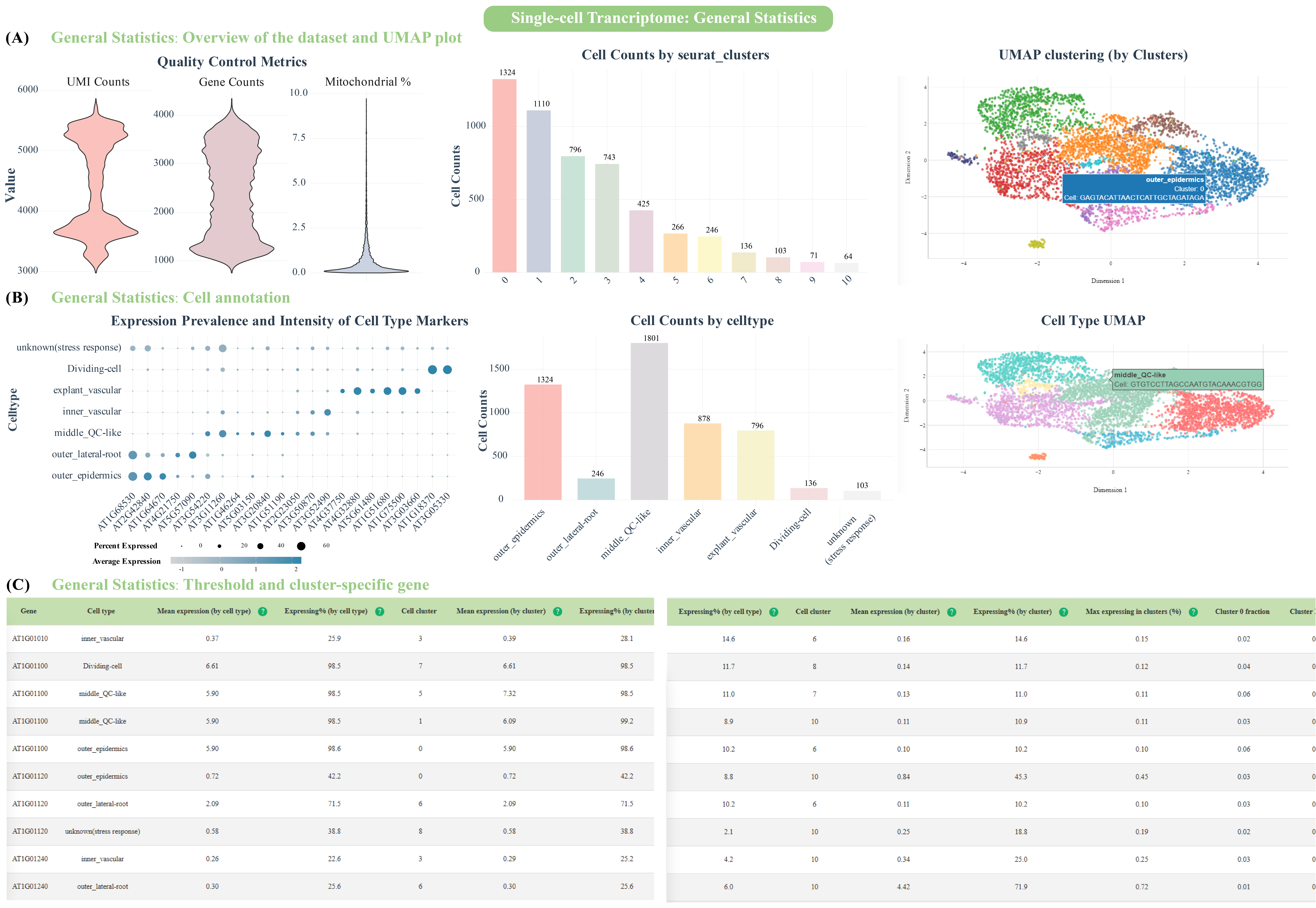

**Figure S10.** **Statistical and analytical functions on the *General Statistics* webpage for single-cell transcriptomic datasets.**

(A) Quality control and dimensionality reduction. Left: violin plots showing distributions of total gene numbers and mitochondrial gene ratios post-filtering; middle: bar plot summarizing cell abundances across 11 clusters; right: UMAP projection of cells colored by corresponding clusters. (B) Cell type annotation. Left: the dot plot illustrates the expression correspondence between known marker genes and annotated cell types. Middle: the bar chart shows the cell count distribution among the seven annotated cell types. Right: UMAP visualization of the PRJNA659737 dataset with seven distinct cell types. (C) Differential expression (left) and cluster‑specific gene (right) analysis.

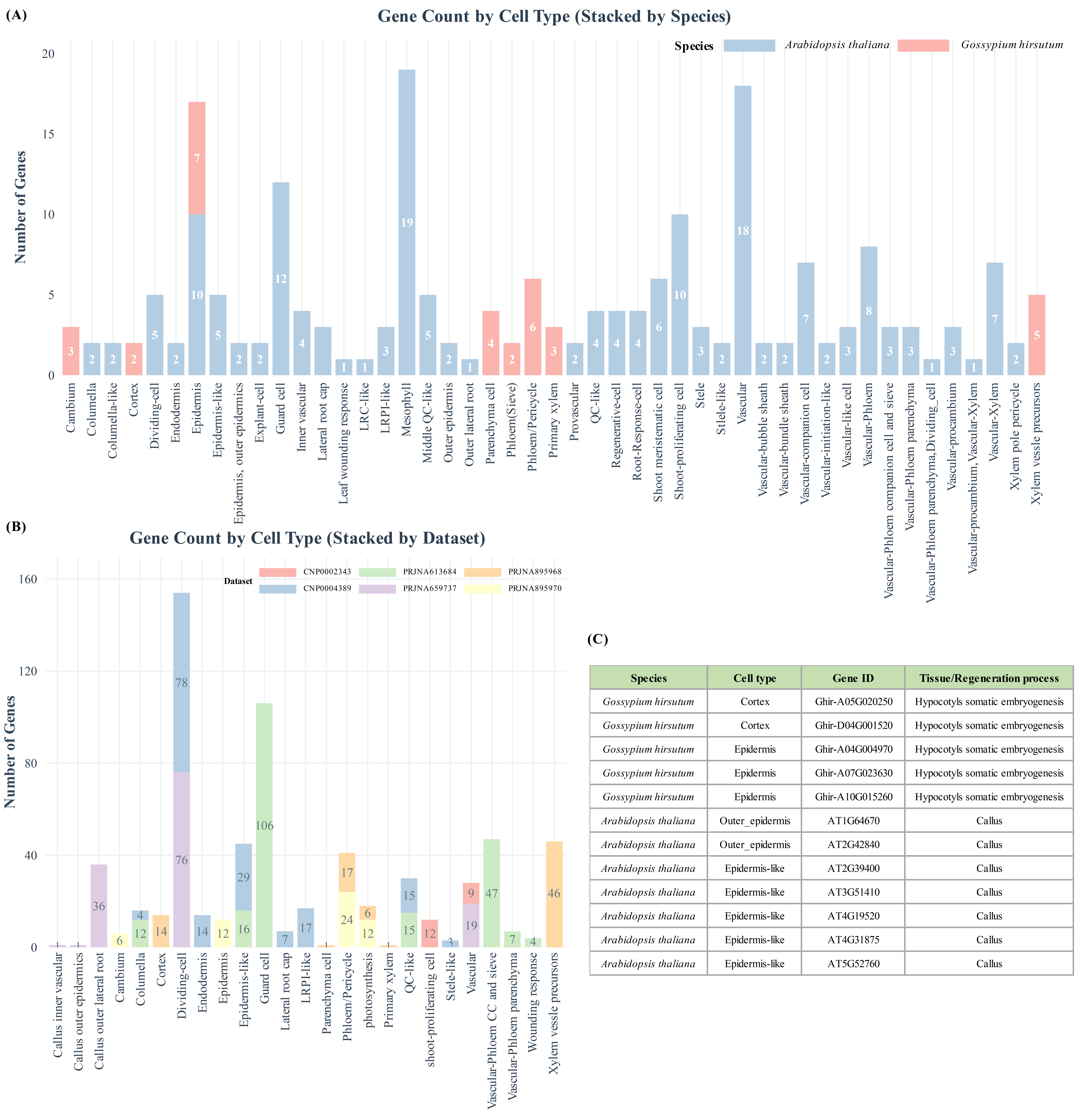

**Figure S11. Curated marker gene collections for cell-type annotation of single-cell datasets.**

(A) Summary statistics of cell types and associated known marker genes. (B) Distribution of candidate marker gene quantities across annotated cell types. (C) Screenshot of the *Marker Genes* webpage showing representative known marker gene information on PlantRegMoD.

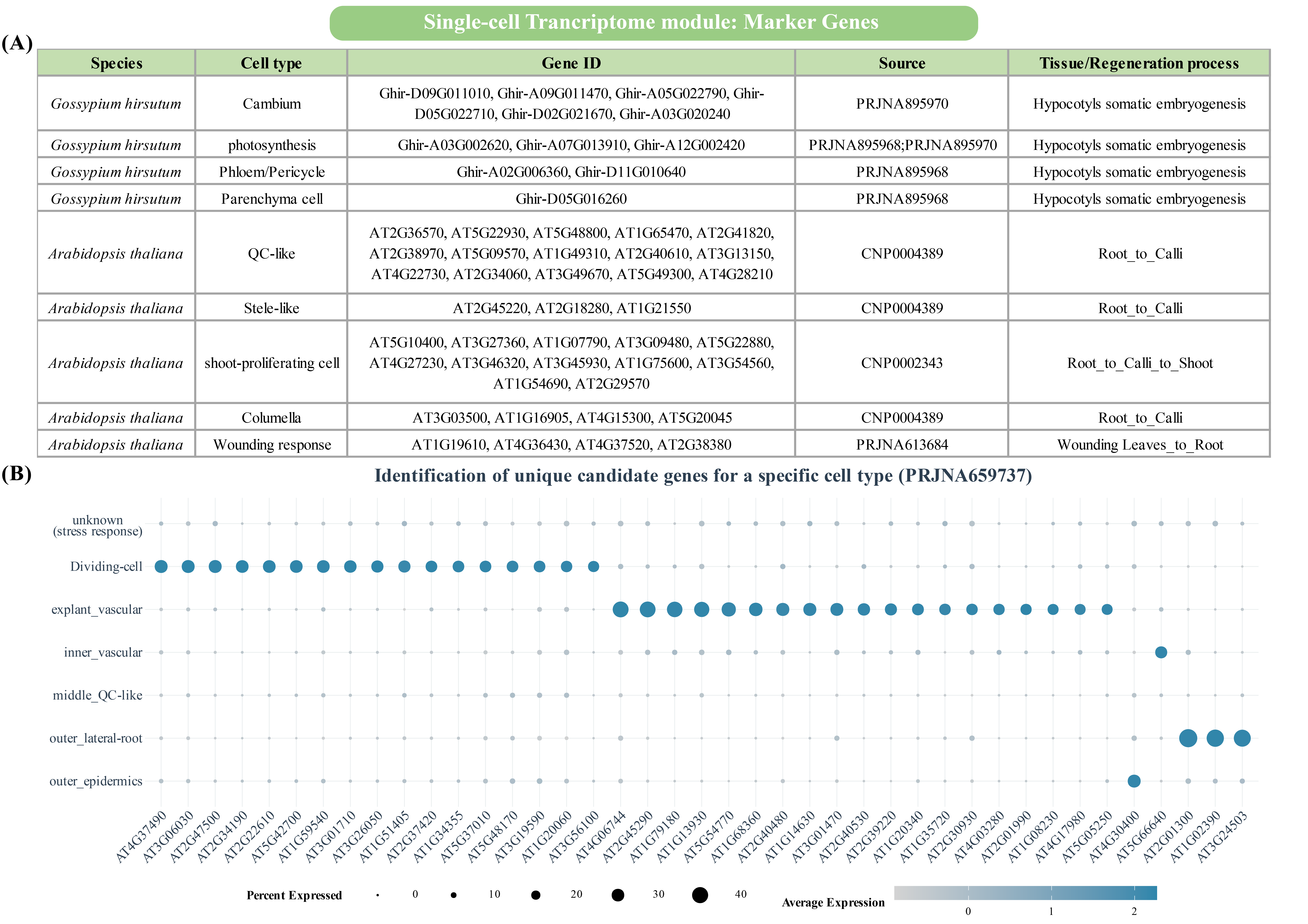

**Figure S12 Candidate marker genes.**

(A) Screenshot showing the specific information of some candidate marker genes on the "Marker Genes" webpage of the database. (B) Dot plot showing expression of some candidate cell-type specific genes.

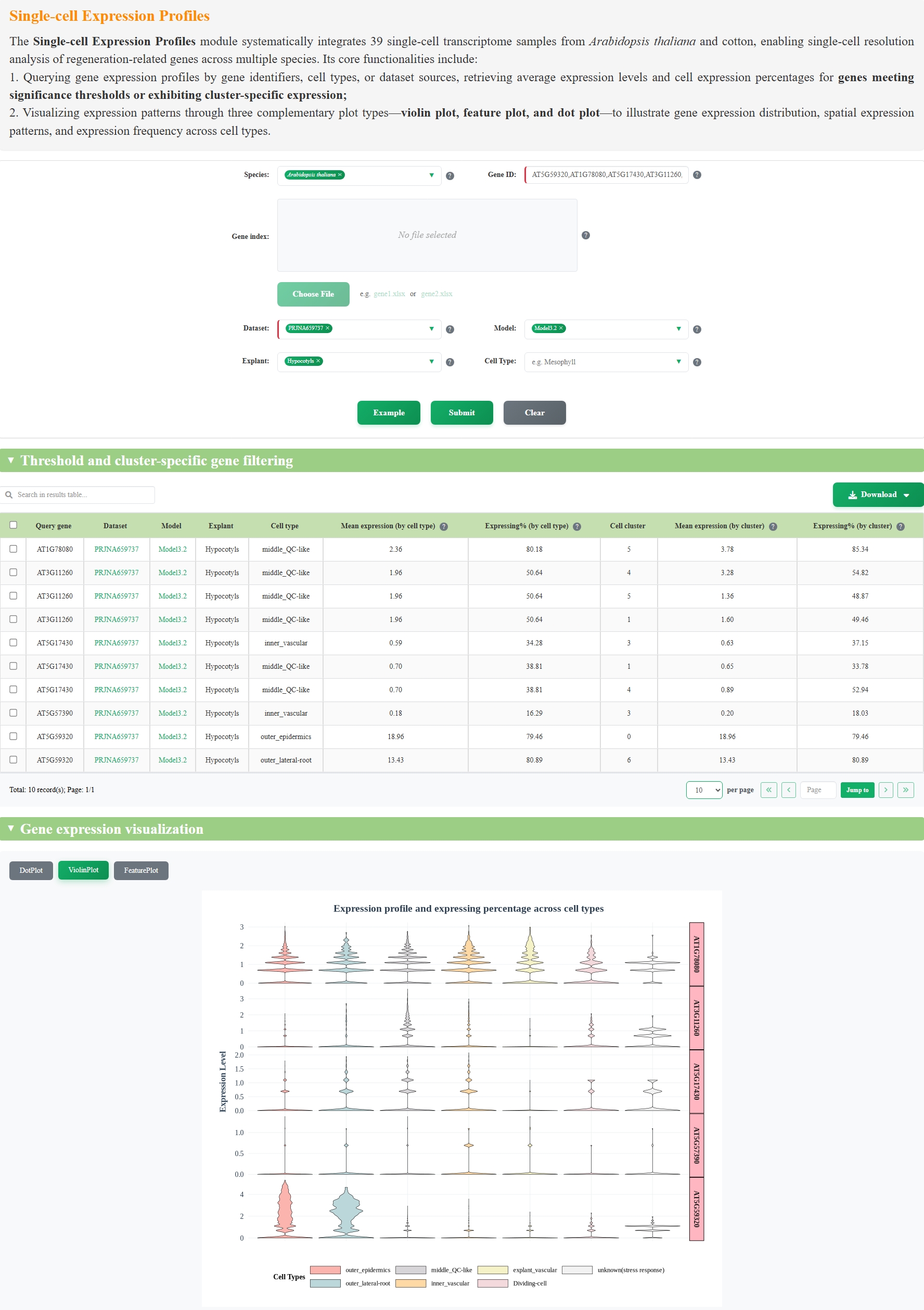

**Figure S13. Screenshot showing single-cell expression profiles of representative genes.**

Tabular statistics and dot-plot visualization of *LTP3* (AT5G59320), *WIND1* (AT1G78080), *BBM* (AT5G17430), *WOX5* (AT3G11260), and *PLT5* (AT5G57390) in the PRJNA659737 dataset on the *Single-cell Expression Profiles* webpage.

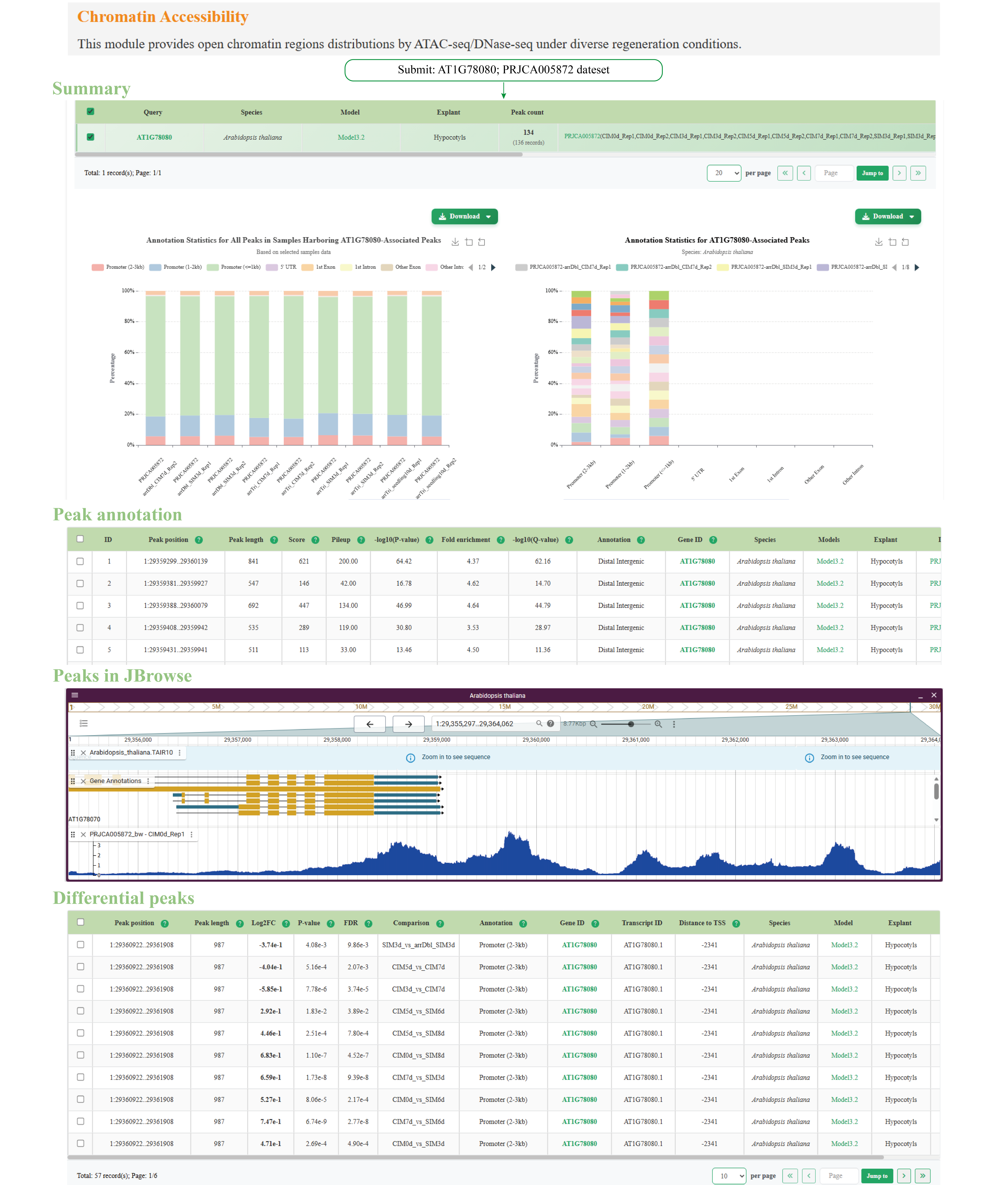

**Figure S14 Example usage of the *Chromatin Accessibility* webpage in the *Epigenome* module.**

Illustrative retrieval of chromatin accessibility peaks surrounding the *Arabidopsis thaliana* *WIND1* gene (*AT1G78080*) across all samples in the PRJCA005872 dataset, with the first entry in the *Peak annotation* table selected for visualization via JBrowse.

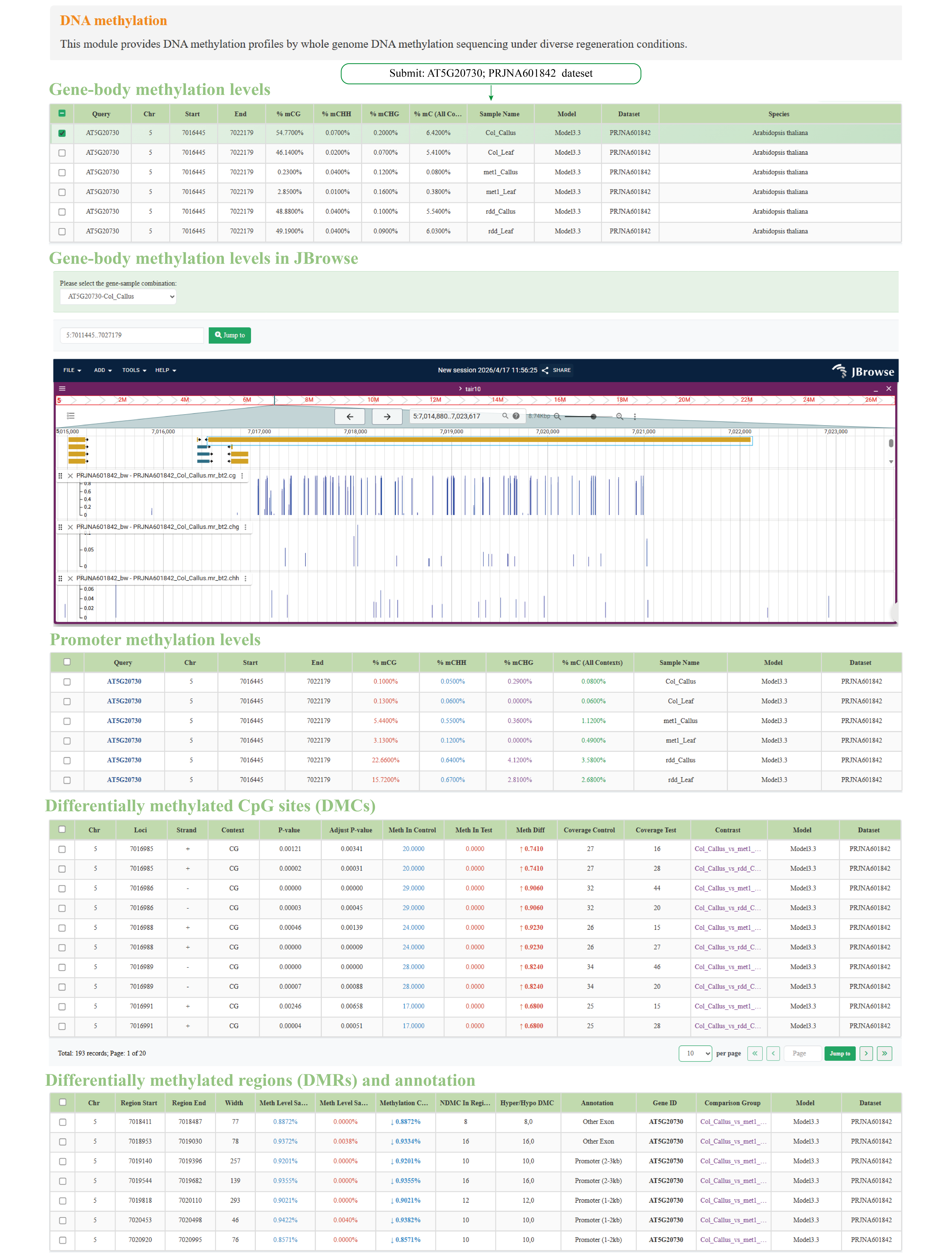

**Figure S15 Example usage of the *DNA* *Methylation* webpage in the *Epigenom*e module.**

Illustrative retrieval of DNA methylation profiles for the *Arabidopsis thaliana* *ARF7* (*AT5G20730*) across all samples in the PRJNA601842 dataset, with the first entry in the *Gene-body methylation levels* and *Promoter methylation levels* tables selected for visualization in JBrowse.

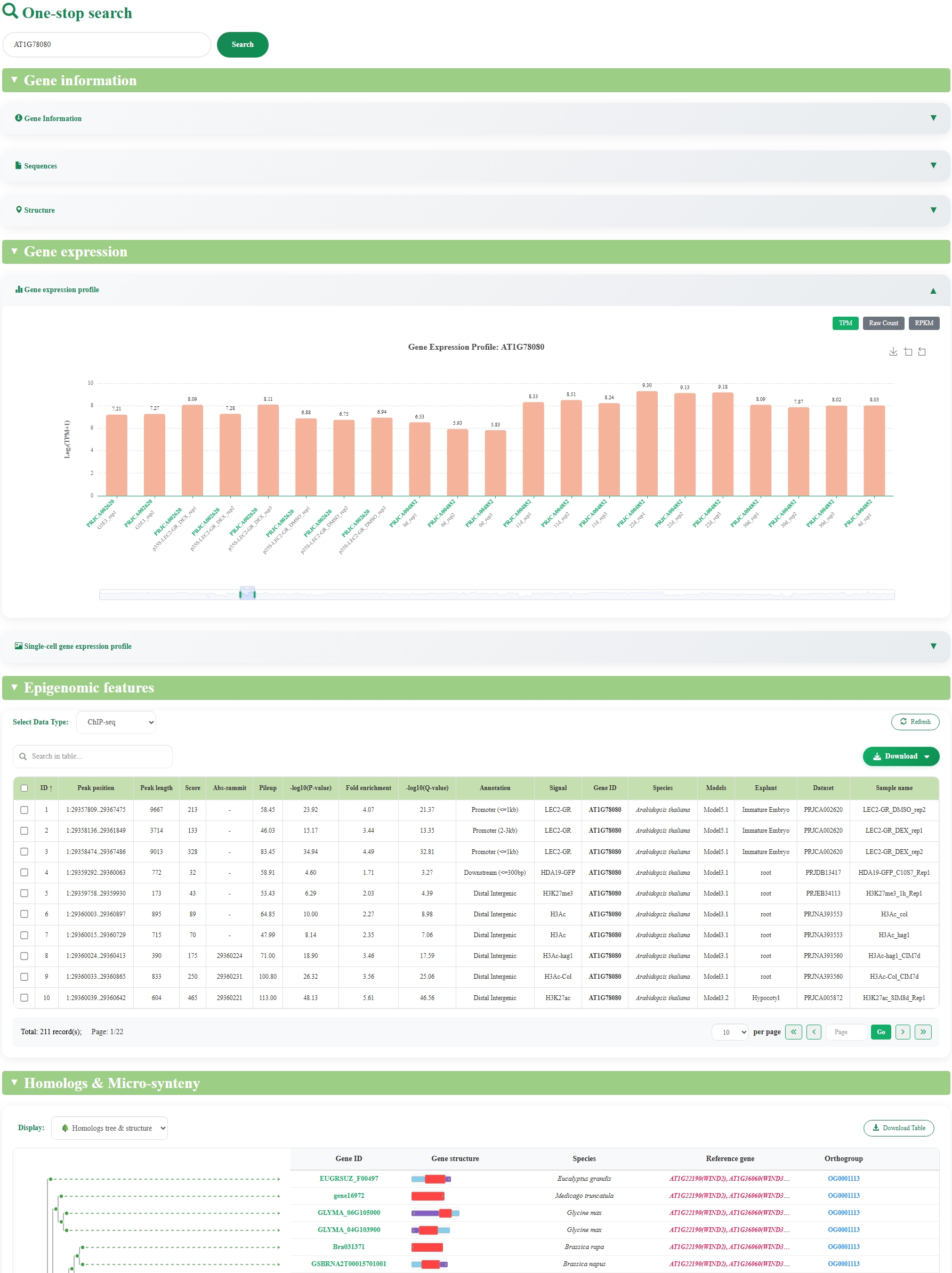

**Figure S16 Screenshot showing an example of one-stop search usage (*WIND1*).**
